# Role of Hypertrophic Adipocytes, Collagen VI and CD38 in Fat Fibrosis of Patients with Obesity

**DOI:** 10.1101/2025.10.08.681115

**Authors:** Angelica Di Vincenzo, Tonia Luca, Jessica Perugini, Giovanni Lezoche, Vincenza Barresi, Vincenzo De Geronimo, Abele Donati, Erika Casarotta, Mario Tomasello, Salvatore Pezzino, Chiara Scuderi, Adriano Di Cristoforo, Alessio Pieroni, Massimiliano Petrelli, Maria Vittoria Napoli, Nayra Figueiredo, Flavia Campos Corgosinho, Andrea Sbarbati, Luciano Merlini, Patrizia Sabatelli, Laura Graciotti, Tatiana Spadoni, Christian Dani, Monica Mattioli-Belmonte, Daniele Condorelli, Armando Gabrielli, Fabio Malavasi, Antonio Giordano, Sergio Castorina, Saverio Cinti

**Affiliations:** Center for the study of Obesity, Department of Experimental and Clinical Medicine, Marche Polytechnic University, Via Tronto 10a, 60126 Ancona, Italy; Department of Medical, Surgical Sciences and Advanced Technologies “G.F. Ingrassia”, University of Catania, Via Santa Sofia, 87, 95123 Catania, Italy; Department of Experimental and Clinical Medicine, Section of Surgical Sciences, Marche Polytechnic University, Via Tronto 10a, 60126 Ancona, Italy; Department of Biomedical and Biotechnological Sciences, Section of Medical Biochemistry, University of Catania, Via Santa Sofia 89, 95123, Catania, Italy; Unit of Endocrinology-Policlinico Morgagni CCD, 95125 Catania, Italy; Anesthesia and Intensive Care Unit, Department of Biomedical Sciences and Public Health, Marche Polytechnic University, Via Tronto 10a, 60126, Ancona, Italy; Mediterranean Foundation “G.B. Morgagni”, Catania, Italy; Department of Medicine and Surgery, Kore University of Enna, Contrada Santa Panasia, Enna, 94100, Italy; Department of Science and Engineering of Matter, Environment and Urban Planning, Via Brecce Bianche 12, 60131 Ancona, Italy; Clinic of Endocrinology and Metabolic Diseases. Marche Polytechnic University. Via Tronto 10a, 60126 Ancona, Italy; Foundation of Molecular Medicine and Cellular Therapy Marche Polytechnic University, Via Tronto 10a, 60126, Ancona, Italy; Postgraduate Program in Health Sciences, Faculty of Medicine, Federal University of Goiás (UFG)- 5th Avenue, s/n - Setor Leste Universitário, Goiânia - GO, 74605-050, Brazil; Department of Neuroscience, biomedicine and movement sciences. Strada le Grazie 8, 37134 Verona, Italy; Department of Biomedical and Neuromotor Science, University of Bologna, DIBINEM, 40136 Bologna, Italy; CNR, Institute of Molecular Genetics “Luigi Luca Cavalli-Sforza”, Via di Barbiano 1/10, 40136 Bologna, Italy; Center of Confocal, Electron Microscopy and CLEM Department of Biomedical Science and Public Health Faculty of Medicine- Marche Polytechnic University Via Tronto, 10 60126 Ancona, Italy; Faculté de Médecine, CNRS, INSERM, iBV, Université Côte d’Azur, CEDEX 2, F-06107 Nice, France; Department of Clinical and Molecular Sciences, Marche Polytechnic University, Via Tronto 10a, 60126, Ancona, Italy; Department of Medical Sciences, University of Torino Medical School, Fondazione Ricerca Molinette Ets, Via Valeggio 41, 10129 Torino, Italy

## Abstract

Fat fibrosis correlates to metabolic consequences in patients with obesity, and is due to three types of collagen: I and III (fibrillar) and VI (non-fibrillar).

In this sudy the extent of fibrosis in obese patients (n 50) was significant only in visceral parenchymal fat (4.7% vs 2.5% in controls (n 15) P<0.0001) and not in subcutaneous fat. Electron microscopy, in vivo and in vitro data, suggested that obese adipocytes are responsible for fibrillar collagen (I and III) production. COL6 (gene producing the non fibrillar form) resulted less expressed. In line, patients with COL6 mutations, showed increased fibrotic tissue even in subcutaneous fat: about 6.5 times vs controls in the patient with the severe form (Ullrich) and 2.8 times in two patients with the milder form (Bethlem).

Approximately 15% of obese adipocytes were dead (perilipin1 negative), and consequent infiltrating macrophages showed hyperexpression of CD38, an ectoenzyme implicated in systemic fibrosis. Correlations with gene expression confirmed the importance also of myofibroblasts and the extracellular matrix peptidase D.

All together our data support a role for obese adipocytes in the fibrillar collagen production and evidentiate collagen VI and CD38 as new molecular determinants, reinforcing the idea of a multi-factorial origin of fat fibrosis.

## Introduction

Adipose tissue is a complex organ, recently recognized as forming a unitary structure in both mice and humans^1^. In individuals with obesity, the adipose organ is marked by chronic low-grade inflammation and fibrosis^2,3^. While the pathogenesis and clinical consequences of chronic inflammation are well understood, the role of fibrosis remains less clear and is still a matter of debate. Nonetheless, its importance is highlighted by studies linking visceral fat fibrosis to glucose and lipid dysmetabolism^4,5^, as well as evidence suggesting potential oncogenic associations^6^.

In obesity, adipose tissue primarily contains three types of collagen (Col): types I, III and VI^7^. While Col I and Col III are mainly associated with the classical mechanical consequences of fibrosis^8,9^, the role of Col VI is much more complex and not fully understood^10–13^.

Although Col VI is expressed throughout the body, it is particularly abundant in the extracellular matrix of adipose tissue^7,14^, suggesting it plays a key role in adipose tissue physiology.

Mutations in Col VI genes (*COL6A1*, *COL6A2* and *COL6A3*) cause a group of myopathies (COLVI- RM) characterized by muscle weakness, joint contractures and skin abnormalities. They represent a continuum of clinical phenotypes from early severe forms (Ullrich Congenital Muscular Dystrophy, UCMD) to milder presentations (Bethlem Myopathy, BM) and intermediate phenotypes^15^.

In this study, we analyzed subcutaneous and visceral (omental) fat biopsies from 50 patients with obesity undergoing bariatric surgery and 15 lean control patients. Morphometric analysis of Sirius Red-stained histological sections revealed significant fibrosis only in visceral fat, consistent with previous findings^3,16,17^.

As expected, gene expression of *COL1A1* and *COL3A1* (most representative isoforms of Col I and III) was elevated in visceral fat, while the expression of *COL6A3* (most representative isoform of Col VI) was reduced.

Transmission electron microscopy (TEM) and high-resolution scanning electron microscopy (HRSEM) showed that fibrillar collagen is a fundamental component of the adipocyte surface anatomy. In obese adipocytes, fibrillar collagen appears to increase, likely in response to decreased Col VI, as suggested by gene expression data and in vitro silencing experiments. Furthermore, fat biopsies from patients carrying homozygous (Ullrich) or heterozygous (Bethlem) mutations of the Col VI genes showed high levels of fibrosis, exceeding those observed in obese adipose tissue. These findings confirm the functional relevance of Col VI in the development of fat fibrosis.

CD38, an ectoenzyme implicated in the pathogenesis of systemic sclerosis, an immune disease characterized by skin and organ fibrosis ^18,19^, also showed a positive correlation with visceral fat fibrosis, accompanied by macrophage-driven inflammation. Immunohistochemical analysis revealed strong CD38 immunoreactivity in both macrophages and obese adipocytes, suggesting that CD38 may contribute to fat fibrosis through an additional molecular pathway. However, gene modulation experiments did not demonstrate a direct functional link between Col VI and CD38, indicating that their roles in fat fibrosis are independent.

Overall, our findings align with previous studies^3,20–22^ in suggesting that fat fibrosis in humans affected by obesity is multifactorial originally, and point to obese adipocytes as a source of collagen production. Moreover, our data highlight an unexpected role for both Col VI and CD38 in driving the altered metabolism of obese visceral cells, ultimately leading to fat fibrosis.

## Patients and Methods

### Patients

Over the past five years, we collected adipose tissue biopsies from the subcutaneous and omental fat of 50 individuals undergoing bariatric surgery for severe obesity (body mass index [BMI] ≥ 35 kg/m^2^) at Marche Polytechnic University, Ancona (n = 33) and at the “G.B. Morgagni” Polyclinic-University of Catania (n = 17). The clinical characteristics of these patients are summarized in Table I. Fat biopsies from the same anatomical sites were also obtained from control subjects (CTRL, n = 15) (BMI ≤ 25 kg/m^2^ with abdominal circumference within the normal range) undergoing cholecystectomy at the “G.B. Morgagni” Polyclinic-University of Catania (Table II).

**Table I.** Clinical data of patients with obesity.

| Name | Age | Sex | BMI | Glycaemia | Diabetes | Total cholesterol (mg/dl) | HDL cholesterol (mg/dl) | Triglycerides (mg/dl) |
| --- | --- | --- | --- | --- | --- | --- | --- | --- |
| 1 | 36 | M | 47.62 | 107 | No | 211 | 64 | 70 |
| 2 | 35 | F | 42.46 | 106 | No | 157 | 51 | 88 |
| 3 | 35 | F | 44.89 | 102 | Yes* | 229 | 60 | 73 |
| 4 | 41 | F | 52.21 | 121 | No | 214 | 35 | 270 |
| 5 | 17 | F | 40.01 | 85 | No | 172 | 53 | 78 |
| 6 | 39 | M | 44.36 | 134 | Yes* | 212 | 43 | 153 |
| 7 | 37 | F | 48.18 | 113 | No | 199 | 38 | 125 |
| 8 | 29 | F | 50.47 | 96 | No | 220 | 48 | 85 |
| 9 | 35 | F | 38.95 | 86 | No | 141 | 40 | 68 |
| 10 | 37 | M | 41.19 | 100 | No | 135 | 47 | 54 |
| 11 | 27 | F | 38.34 | 97 | No | 228 | 42 | 172 |
| 12 | 42 | M | 41.29 | 104 | Yes* | 194 | 50 | 151 |
| 13 | 34 | F | 41.90 | 82 | No | 173 | 72 | 59 |
| 14 | 21 | F | 46.95 | 89 | No | 163 | 46 | 123 |
| 15 | 47 | F | 56.95 | 106 | Yes* | 254 | 46 | 259 |
| 16 | 26 | M | 41.45 | 63 | No | 173 | 46 | 76 |
| 17 | 44 | M | 37.09 | 102 | No | 241 | 57 | 77 |
| 18 | 56 | M | 44.5 | 161 | No | 137 | 41 | 152 |
| 19 | 57 | F | 30 | 86 | No | 183 | 64 | 60 |
| 20 | 42 | M | 43 | 85 | No | 175 | 42 | 96 |
| 21 | 53 | F | 43.25 | 92 | No | 159 | 49 | 127 |
| 22 | 44 | M | 45.6 | 80 | No | 152 | 47 | 105 |
| 23 | 32 | F | 48 | 85 | No | 165 | 48 | 133 |
| 24 | 38 | F | 50 | 259 | Yes* | 228 | 72 | 234 |
| 25 | 56 | F | 43.9 | 86 | No | / | / | / |
| 26 | 46 | F | 42 | 116 | No | 193 | 51 | 169 |
| 27 | 32 | F | 32 | 71 | No | / | / | / |
| 28 | 48 | M | 38.9 | 78 | No | 167 | 23 | 276 |
| 29 | 41 | F | 40 | 104 | No | 209 | 34 | 191 |
| 30 | 44 | F | 46 | 90 | Yes* | 236 | 45 | 176 |
| 31 | 44 | F | 40.8 | 127 | No | 172 | 36 | 108 |
| 32 | 42 | F | 38 | 74 | No | 225 | 39 | 212 |
| 33 | 46 | F | 40 | 86 | No | 215 | 64 | 133 |
| 34 | 46 | F | 45 | 335 | Yes* | / | / | / |
| 35 | 42 | F | 50 | 104 | No | 258 | 36 | 262 |
| 36 | 54 | F | 43.6 | 224 | Yes* | / | / | / |
| 37 | 55 | F | 43.3 | 79 | No | 238 | 65 | 121 |
| 38 | 56 | F | 41 | 123 | Yes* | 207 | 59 | 118 |
| 39 | 43 | M | 43 | 107 | No | 238 | 46 | 111 |
| 40 | 30 | M | 60 | 86 | No | 158 | 35 | 115 |
| 41 | 45 | M | 41 | 110 | Yes* | 140 | 40 | 111 |
| 42 | 33 | M | 43.3 | 98 | No | 169 | 57 | 57 |
| 43 | 47 | F | 29.76 | 83 | No | / | / | / |
| 44 | 46 | F | 48.28 | 122 | No | 194 | 67 | 89 |
| 45 | 45 | M | 40.9 | 79 | Yes* | 197 | 40 | 137 |
| 46 | 52 | F | 42.7 | 78 | No | / | / | / |
| 47 | 42 | F | 40 | 137 | Yes* | 129 | 32 | 254 |
| 48 | 55 | M | 59.1 | 117 | Yes* | 149 | 42 | 128 |
| 49 | 26 | F | 36.4 | 86 | No | 127 | 36 | 68 |
| 50 | 27 | M | 41 | 87 | No | / | / | / |
| Average±S.E.M. | 40.54±1.37 |  | 43.37±0.86 | 108.6±6.7 |  | 189.2±5.57 | 47.63±1.74 | 133.1±9.73 |

\*Diagnostic criteria by the American Diabetes Association (ADA) include the following:
- fasting plasma glucose (FPG) levels $\geq 125$ mg/dL; - 2-hour plasma glucose $\geq 200$ mg/dL during an Oral Glucose Tolerance Test (OGTT). The test should be performed as described by the World Health Organization (WHO), using a glucose load containing the equivalent of 75 g anhydrous glucose dissolved in water; - glycated haemoglobin $> 6.5\%$ . - glycaemic levels $\geq 200$ mg/dL in the presence of symptoms

**Table II.** Clinical data of lean patients.

| Name | Age | Sex | BMI | Abdominal circumference |
| --- | --- | --- | --- | --- |
| 1 | 59 | M | 23 | 92 |
| 2 | 72 | M | 24 | 90 |
| 3 | 55 | F | 19 | 72 |
| 4 | 49 | F | 24.77 | 83 |
| 5 | 55 | M | 27.3 | 71 |
| 6 | 66 | F | 24.24 | 81 |
| 7 | 47 | F | 20.76 | 70 |
| 8 | 39 | M | 25.08 | 100 |
| 9 | 31 | F | 18.75 | 68 |
| 10 | 43 | F | 18.8 | 73 |
| 11 | 17 | F | 24.8 | 71 |
| 12 | 29 | M | 24.6 | 93 |
| 13 | 23 | M | 24.6 | 91 |
| 14 | 38 | M | 25.5 | 84 |
| 15 | 39 | F | 19.61 | 72 |
| Average±S.E.M. | 44.13±4.04 |  | 22.99±0.73 | 80.73±2.70 |

Subcutaneous fat biopsies were also collected from individuals with Ullrich congenital muscular dystrophy (UCMD) (n = 1) and Bethlem myopathy (BM) (n = 2) at the Rizzoli Orthopedic Institute, Department of Biomedical and Neuromotor Science, University of Bologna. Their clinical and genetic data are presented in Table III.

**TAB III.**
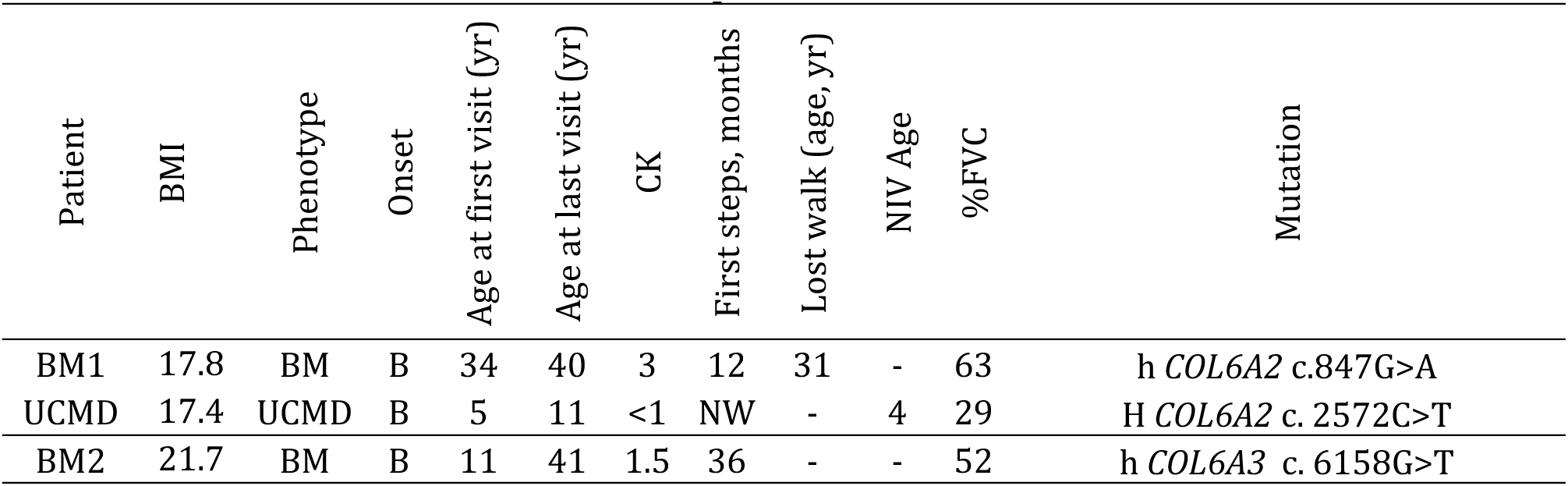
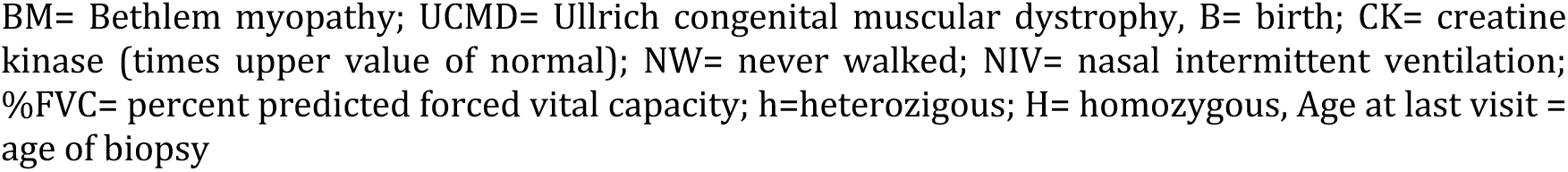
Clinical data of Ullrich and Bethlem patients.

| Patient | BMI | Phenotype | Onset | Age at first visit (yr) | Age at last visit (yr) | CK | First steps, months | Lost walk (age, yr) | NIV Age | %FVC | Mutation |
| --- | --- | --- | --- | --- | --- | --- | --- | --- | --- | --- | --- |
| BM1 | 17.8 | BM | B | 34 | 40 | 3 | 12 | 31 | - | 63 | h *COL6A2* c.847G>A |
| UCMD | 17.4 | UCMD | B | 5 | 11 | <1 | NW | - | 4 | 29 | H *COL6A2* c. 2572C>T |
| BM2 | 21.7 | BM | B | 11 | 41 | 1.5 | 36 | - | - | 52 | h *COL6A3* c. 6158G>T |

BM= Bethlem myopathy; UCMD= Ullrich congenital muscular dystrophy, B= birth; CK= creatine kinase (times upper value of normal); NW= never walked; NIV= nasal intermittent ventilation; %FVC= percent predicted forced vital capacity; h=heterozygous; H= homozygous, Age at last visit = age of biopsy

Additionally, subcutaneous calcaneal fat samples from four patients aged 27-47 years were collected at Verona University, Department of Neuroscience, Biomedicine and Movement Sciences, treated for retrocalcaneal exostosis.

### Light microscopy

For morphological analysis by light microscopy, all omental and subcutaneous fat samples were fixed in 4% paraformaldehyde in 0.1 M phosphate buffer (pH 7.4) overnight at 4 ° C. The samples were then dehydrated through a series of alcohol solutions of increasing concentration, clarified in xylene, and embedded in paraffin blocks. Serial paraffin sections (3 μm thick) were cut from each adipose biopsy using a sliding microtome (Leica RM 2135, Leica Microsystem, Milan, Italy), mounted onto glass slides, and dried.

### Histochemistry with morphometry

Fibrosis quantification was performed using Sirius Red staining^23^.

Paraffin sections (3μm thick) were dewaxed in xylene and rehydrated through sequential immersions in alcohol solutions of decreasing concentration. The sections were then incubated with Picro-Sirius Red dye for one hour at room temperature. After abundant washing in water to remove excess dye, the sections were contrasted with Hematoxylin for 2 min, rinsed, dehydrated, and mounted using Eukitt® Mounting Medium (Merk Life Science S.r.l., Milan, Italy). Images of Sirius Red-stained tissues were captured at 2.5x magnification, with an average of 20 images acquired per biopsy. Fibrosis was semi-automatically quantified using ImageJ software: images were converted to grayscale, the red-stained areas were selected using the threshold command, and the percentage of total fibrosis was calculated. Fibrosis was expressed as the ratio of the red-stained area (fibrotic) to the total tissue area examined.

### Immunohistochemistry with morphometry

Paraffin sections (3 μm thick) were rehydrated deparaffinized and rehydrated. Unmasking procedures, with citric acid at 96 °C for 20 min, were used for the immunohistochemical detection of CD68. Then, sections were reacted with 0.3% hydrogen peroxide solution in distilled water for 5 min to block endogenous peroxidase activity. After rinsing in PBS, sections were treated for 20 min with a 2% normal serum blocking solution in PBS to block non-specific binding of the secondary antibody. The sections were then incubated overnight at 4°C with the following primary antibodies: polyclonal rabbit anti-perilipin 1 (ab3526) 1:800 in PBS (Abcam, Cambridge, England); monoclonal mouse anti-CD68 (M0814) 1:200 in PBS (Dako, Glostrup, Denmark) and monoclonal antibody murine anti-human CD38, clone IB4, IgG2a Class, 1:300 in PBS^24^.

After rinsing in PBS, biotinylated secondary antibody was added (1:200 dilution in PBS for 30 min; Vector Laboratories, Burlingame, CA). Sections were rinsed again in PBS and incubated with Avidin-Biotin peroxidase complex (1:100 in PBS for 60 min; Vectastain ABC kit Vector Laboratories). Sections were rinsed in PBS several times and finally the substrate Sigma Fast 3,3-diaminobenzidine (Sigma–Aldrich, Vienna, Austria) was added for 3 min. After immunohistochemical staining, sections were counterstained with Hematoxylin, dehydrated in ethanol, cleared with xylene, and mounted with Eukitt® Mounting Medium (Merk Life Science S.r.l., Milan, Italy). Negative controls, in which the primary antibody was omitted, showed no staining.

Morphometric analyses were performed on tissue sections observed with a light microscope (Zeiss Axioscop 40) (Carl Zeiss GmbH, Jena, Germany) to measure adipocyte size, the immunoreactive areas composed of perilipin 1 (PLIN1)-positive (vital) adipocytes, and the inflammatory state. All images were captured with a Zeiss Axiocam 503 color digital camera.

Adipocyte size: images were captured at 10x magnification and the mean area of at least 200 PLIN1 immunoreactive adipocytes per biopsy was calculated using ImageJ software.

Viability of adipocytes: sections were captured at 2.5x magnification to acquire a number of images corresponding approximately to the full size of the tissue section. Using ImageJ software, the total adipose tissue area composed of PLIN1-positive and PLIN1-negative adipocytes was measured. The percentage of PLIN1 negativity was then calculated for each sample.

Inflammatory state: quantification of CD68 positive cells was performed. Approximately 20 random fields per tissue section were captured at 20x magnification, and CD68-positive macrophages were counted using ImageJ software. Counts were normalized to 10^4^ adipocytes. Macrophages were categorized based on their distribution in five tissue districts: around dead adipocytes (arranged in crown-like-structures: CLS)^25^; within fibrotic areas; within the parenchyma; inside blood vessels (intravascular); near blood vessels (perivascular). For each patient, the percentage of CD68-positive macrophages in each district was calculated. Additionally, CLS density (number of CLS per 10^4^ adipocytes) was determined.

### Transmission Electron Microscopy (TEM)

Some of the biopsies (24 in total, 18 from patients with obesity and 6 from lean patients) were also used for electron microscopy studies. For ultrastructural analysis, biopsies were fragmented into pieces measuring approximately 1 mm^3^. Small tissue fragments were fixed in 2% glutaraldehyde and 2% paraformaldehyde in 0.1 mol/L Phosphate Buffer (PB), pH 7.4 for 4 h at room temperature. Samples were then post-fixed in 1% osmium tetroxide in 0.1 mol/L PB (1 h at 4°C), dehydrated through graded acetone series, and embedded in a mixture of Epon-Araldite. For each sample, semi-thin sections were cut (1,5 µm thick), and stained with toluidine blue. Ultrathin sections (60 nm) were then obtained using an MT-X ultratome (RMC; Tucson, AZ), stained with lead citrate, and examined with a Philips CM 10 transmission electron microscope (Philips, Eindhoven, Netherlands) operating at 100kV. For each patient, at least three different thin sections, selected based on semithin toluidine blue staining, were observed. Quantitative data on fibril size were obtained through direct measurements during observations, with at least 40-50 measurements per sample with a soft imaging system GmbH (Muenster, Germany) coupled with MegaView G2 TEM camera (Evident, Waltham, MA, U.S.), performed per sample at high magnification, corresponding to the magnification shown in Fig 2.

### High-Resolution Scanning Electron Microscopy (HRSEM)

Small fragments of omental (n = 6) and subcutaneous (n = 6) adipose tissue from 6 patients affected by obesity and with fibrosis >5% and 3 lean subjects were fixed in 2% glutaraldehyde– paraformaldehyde overnight at 4°C. Samples were post-fixed in 1% % osmium tetroxide (OsO4) in PB for 60 min at 4°C, washed in PB, and dehydrated through increasing ethanol concentrations. Dehydration was completed by Critical Point Drying, consisting of incubation in increasing concentration of hexamethyldisilazane (HMDS), until embedding in 100% HMDS. Samples were mounted on aluminum stubs using cell adhesive carbon disks and sputter-coated with gold particles prior to SEM observation with a FESEM ZEISS SUPRA 40 microscope (Carl Zeiss NTS GmbH, Oberkochen, Germany). This FESEM has a theoretical resolution of 2 nm when operating at 10 kV and with a working distance of 2 mm.

Quantitative data on fibril size were obtained by direct measurements during observations, with at least 40-50 measurements performed per sample at high magnification, corresponding to the magnification shown in Fig 2.

### Intra-abdominal pressure measurements

We prospectively collected data from adult male patients admitted for brain injury to a single ICU (Anesthesia and Intensive Care Unit, University Hospital “Azienda Ospedaliero-Universitaria delle Marche”, Ancona, Italy). Exclusion criteria included recent abdominal surgery, pancreatitis, cirrhosis with ascites, peritonitis or other abdominal infections, intra-abdominal or retroperitoneal masses, abdominal bleeding, retroperitoneal hematoma, and mechanical ventilation with PEEP values higher than 10 cmH2O.

After verifying all inclusion and exclusion criteria, intra-abdominal pressure (IAP) was measured using the Foley manometer technique. The manometer was placed between the catheter and the drainage device and primed with 20 ml of sterile saline solution. The “0 mmHg” reference mark was aligned with the symphysis pubis, and the filter was elevated vertically above the patient to record the bladder pressure, considered a surrogate measure of intra-abdominal pressure. Measurements were performed within the first 24 hours after ICU admission.

Demographic and anthropometric data were collected, and waist circumferences were measured using abdomen CT scans. Patients were categorized into two groups: those with abdominal (visceral) obesity (waist circumference > 102 cm, n 12) and those without (waist circumference < 102 cm, n 12). All data were anonymized and analyzed in a blind fashion.

### Molecular biology

#### Human samples

Fat biopsies were also used for molecular studies. Each biopsy was snap-frozen in liquid nitrogen, stored at −80 °C and used oppure for determination of gene expression.

#### Cell culture and treatments

Human multipotent adipose-derived stem cells (hMADS), kindly provided by Prof. Christian Dani (Université Côte D’Azur, Nice, France) were cultured as described previously^26^. In brief, hMADS grown in low-glucose (1 g/L) proliferation medium (Dulbecco’s Modified Eagle’s Medium [DMEM]) (Pan-Biotech GmbH (Aidenbach, Germany) supplemented with 10% FBS (Pan-Biotech GmbH (Aidenbach, Germany) and 2.5 ng/ml hFGF-2 (human recombinant fibroblast growth factor, PeproTech, London, UK) were used between the 16th and the 19th passage. To induce adipose differentiation, they were seeded in proliferation medium on multi- well plates at a density of 4500 cells/cm^2^. Upon reaching confluence, hFGF-2 was not replaced. The next day (designated day 0), cells were incubated in an adipogenic medium (serum-free proliferation medium/Ham’s F-12 medium) supplemented with: 10 μg/ml transferrin, 5 μg/ml insulin, 0.2 nM triiodothyronine, 100 μM 3-isobutyl-1-methylxanthine, 1 μM dexamethasone, and 100 nM rosiglitazone. Dexamethasone and 3-isobutyl-1-methylxanthine were discontinued from day 3, and rosiglitazone from day 9. Treatments and biological assays were carried out on differentiated hMADS from day 12 to day 15.

Afterward, hypertrophic-like cells were obtained by treating differentiated hMADS for 12 days with adipogenic medium supplemented with fatty acids i.e., 350 μM palmitate and 350 μM oleate (Sigma–Aldrich, Milan, Italy). This concentration reflects the pathological levels of fatty acids (200–375 μM) adipose cells of obese individuals are exposed to. Fatty acid mixtures were replenished every 3 days, and hypertrophic-like adipocytes were obtained by day 32^27^. For morphometric analyses of lipid droplets (LDs), images were captured at 10x magnification and the mean area of at least 100 LDs in differentiated hMADS and in differentiated hypertrophic hMADS was calculated using ImageJ software.

We also used a human dermal fibroblast cell line, NhDF^28^ (Thermo Fischer Scientific, USA), cultured in DMEM with 10% FBS.

#### *COL6A3* gene silencing and modulation of CD38 gene expression

For COL6A3 gene silencing, Lipofectamine® RNAiMAX (Invitrogen, Carlsbad, USA) was used to transfect differentiated hMADS according to the manufacturer’s reverse transfection protocol. Briefly, siRNAs were diluted in OPTI-MEM medium (siRNA final concentration was 2 µM) with Lipofectamine. Differentiated hMADS were transfected with siRNA *COL6A3* (siRNA ID: s3317) for silencing, while control cells were transfected with non-targeting negative control siRNA (scramble sequence) (both from Thermo Fisher Scientific, USA). Cells were incubated for 4 or 7 days.

For modulation of CD38 gene expression, hMADS, were incubated for 48 hours with 1 μM trans-Retinoic Acid (Sigma-Aldrich, USA).

COL1A1, COL3A1, COL6A3 gene expression, as well as CD38 gene modulation were assessed by qRT- PCR.

#### qRT-PCR

Total RNA from fat biopsies or from hMADS was extracted with TRIZOL reagent (Invitrogen, Carlsbad, CA), purified, digested with ribonuclease-free deoxyribonuclease, and concentrated using the RNeasy Micro Kit (Qiagen, Milan, Italy) according to the manufacturer’s instructions. RNA concentrations were quantified using Nanodrop 1000 (Thermo Fisher Scientific, USA). For determination of mRNA levels, 1 μg RNA was reverse transcribed with a High-Capacity cDNA RT Kit with RNase Inhibitor (Applied BioSystems, Foster City, CA) in a total volume of 20 μl. qRT-PCR was performed using TaqMan Gene Expression Assays and Master Mix TaqMan (Applied Biosystems). All probes were from Applied Biosystems (Suppl. Table IA). Reactions were carried out in a Step One Plus Real-Time PCR system (Applied BioSystems) using 50 ng cDNA in a final reaction volume of 10 μl. The thermal cycle protocol included initial incubation at 95 °C for 10 min followed by 40 cycles of 95 °C for 15 s and of 60 °C for 20 s. A control reaction without reverse transcriptase in the amplification mixture was performed for each sample to rule out genomic contamination. Samples not containing the template were included in all experiments as negative controls. TATA box-binding protein was used as an endogenous control to normalize gene expression. Relative mRNA expression was determined using the ΔCt method (2^−ΔCt^).

For other qRT-PCR analyses, a StepOne TM Real-Time PCR System and a Power SYBR™ Green PCR Master Mix (Applied Biosystems, Foster City, CA, USA) were used, according to the manufacturer’s protocol and as previously reported^29,30^. Each sample was analyzed in triplicate and the average was normalized to human ACTB expression.

Primers were designed by the “Primers-BLAST” tool from NCBI (https://www.ncbi.nlm.nih.gov/tools/primer-blast accessed on June 2023); transcript forward and reverse primer sequence, annealing temperature and fragment size are reported in Suppl Table IB.

**Suppl TABLE IA.**
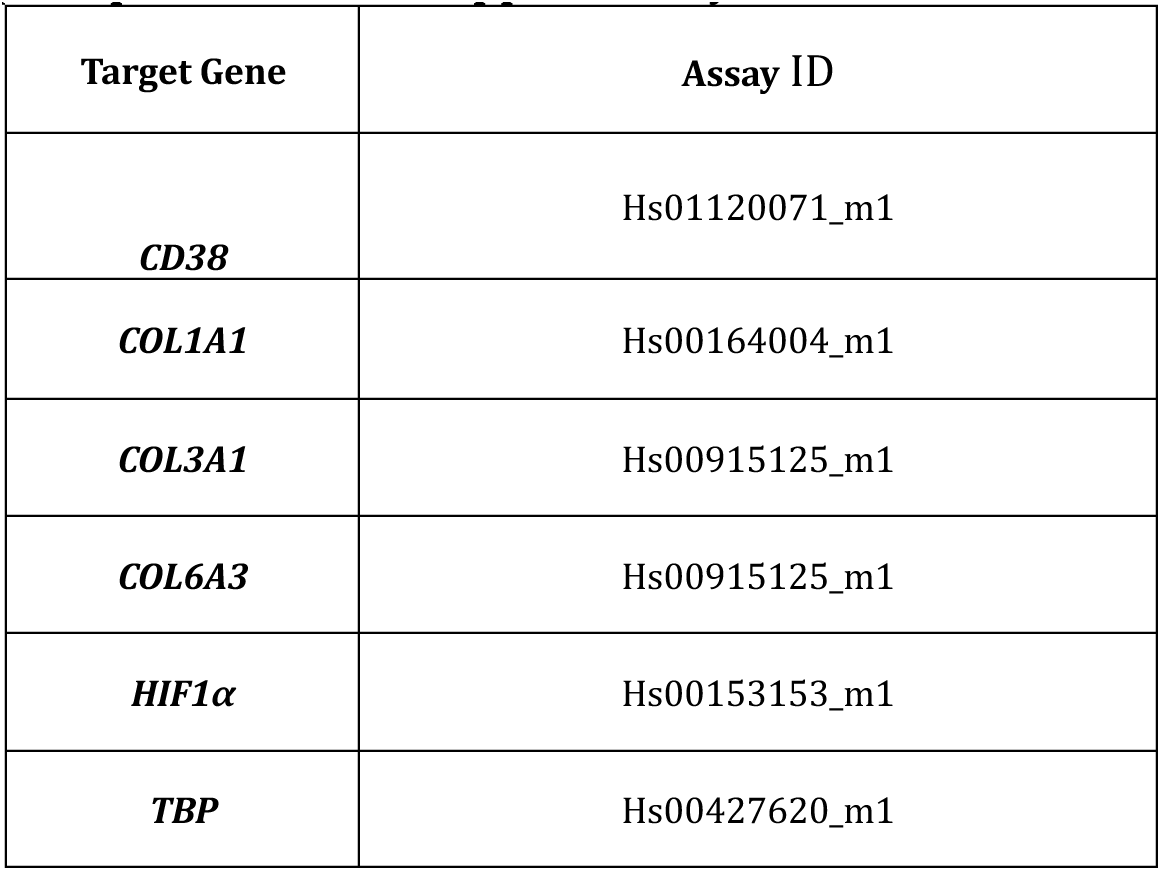
Taqman probes all from Applied Biosystems #4453320.

| Target Gene | Assay ID |
| --- | --- |
| *CD38* | Hs01120071_m1 |
| *COL1A1* | Hs00164004_m1 |
| *COL3A1* | Hs00915125_m1 |
| *COL6A3* | Hs00915125_m1 |
| *HIF1α* | Hs00153153_m1 |
| *TBP* | Hs00427620_m1 |

**Suppl TABLE IB.**
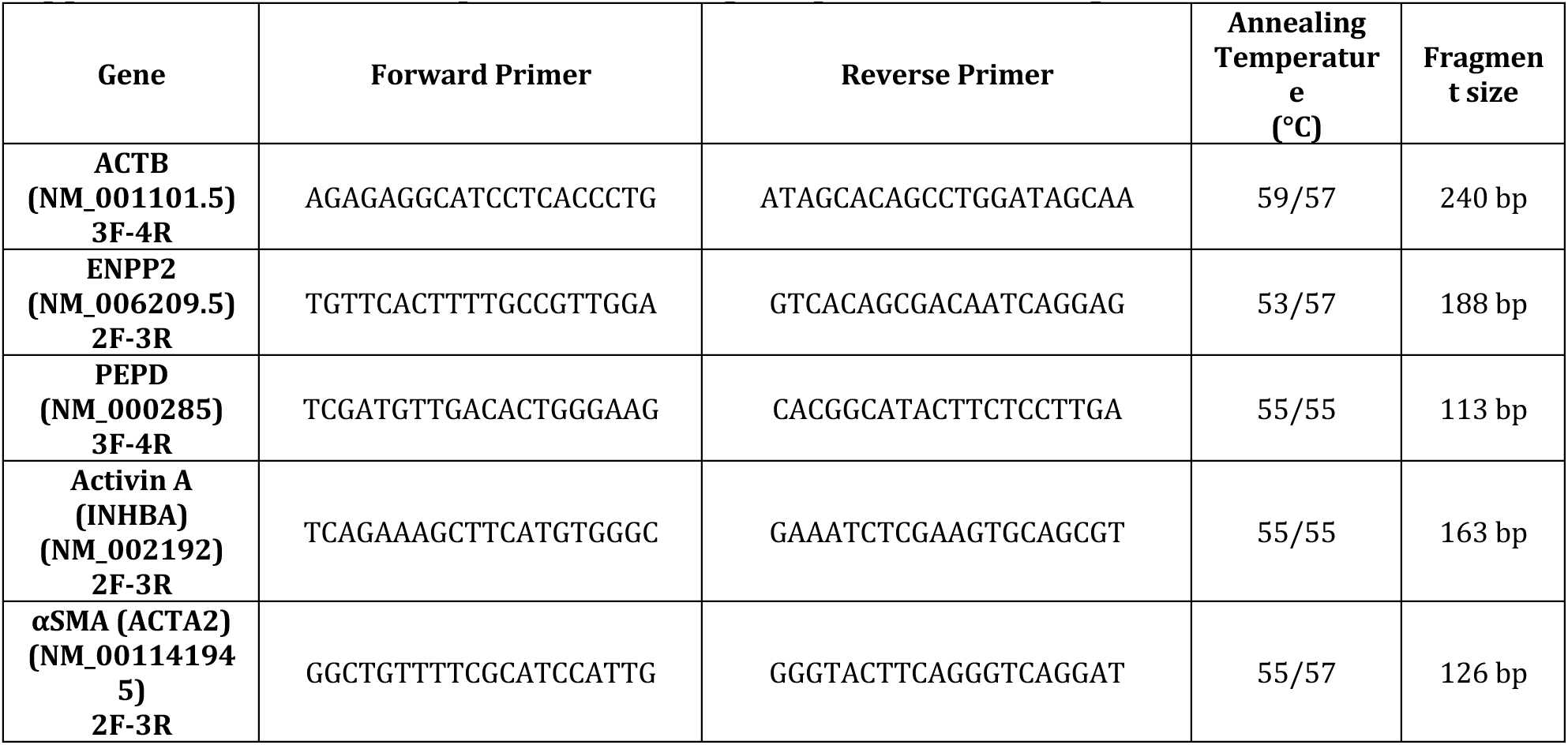
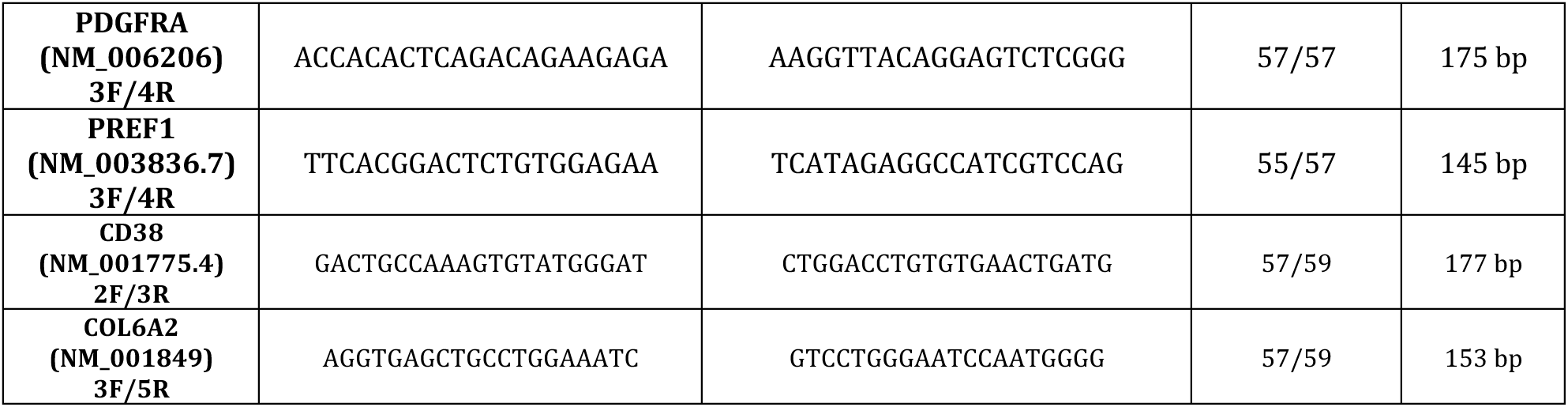
Primer, sequence annealing temperature and fragment size.

Amplification conditions included a cycle at 95 °C for 10 min, followed by 40 cycles at 95 °C for 15 s and 55–59 °C for 1 min. As a negative control, reaction without cDNA was performed (no template control, NTC). The RNA expression level for each sample was calculated using ΔCt values, normalized versus ACTB and reported as 2-^ΔΔCt^.

### Statistical Analyses

All experimental data are presented as mean ± S.E.M. Comparisons between groups (e.g., fibrosis, negativity to PLIN1, CD68 positivity, mean adipocyte size, and gene expression) in lean patients and patients affected by obesity were performed using non-parametric Mann−Whitney or Kruskall-Wallis tests.

For in vitro experiments, one-way analysis of variance (ANOVA) or two-tailed Student’s t-test was applied. Finally, Pearson correlation coefficients (r) and p-values were calculated to assess correlations between variables. Differences were considered significant at p ≤ 0.05.

### Ethical Approval

The study was conducted in accordance with the Declaration of Helsinki and was approved by the following ethics committees: Ancona (CERM 26/11/2020 Prot n. 2020-385 and the study protocol for intra-abdominal pressure measurements -protocol n. 2021 166); Catania (Research Ethics Committee Catania2, Garibaldi Hospital, 21/05/2021 Prot n. 374/CE); and Bologna (11/05/2021, project identification code: CE 0007151). All patients provided written informed consent.

Regarding samples collected at Verona University, patients consented to the anonymous use of their tissues, which would otherwise have been discarded. The materials were enumerated to maintain consistency across the different centers in which they were collected, but the samples were completely anonymized, and no link remains between the material and the patient, in accordance with current regulations (https://english.ccmo.nl/investigators/legal-framework-for-medical-scientific-research/your-research-is-it-subject-to-the-wmo-or-not;^31,32^.

Therefore, no additional ethical approval was required.

## Results

### Omental fat of patients affected by obesity shows parenchymal fibrosis

Adipocytes from subcutaneous and visceral (omental) fat in patients suffering from obesity are widely considered hypertrophic^33^, and our quantitative data fully confirm that both visceral and subcutaneous adipocytes are enlarged compared to those of lean patients, with greater hypertrophy observed in subcutaneous than in visceral fat (Suppl Fig 1). This finding is consistent with previous data^34^. Notably, we measured only perilipin1 (PLIN1) positive adipocytes to exclude confounding results from dead or unhealthy adipocytes (see below).

Sirius Red staining and high-resolution scanning electron microscopy (HRSEM) revealed a variable amount of fibrosis, both surrounding groups of adipocytes and encasing single adipocytes (pericellular fibrosis), in line with data from other studies^3,35^ (Fig 1A).

**Fig 1.**
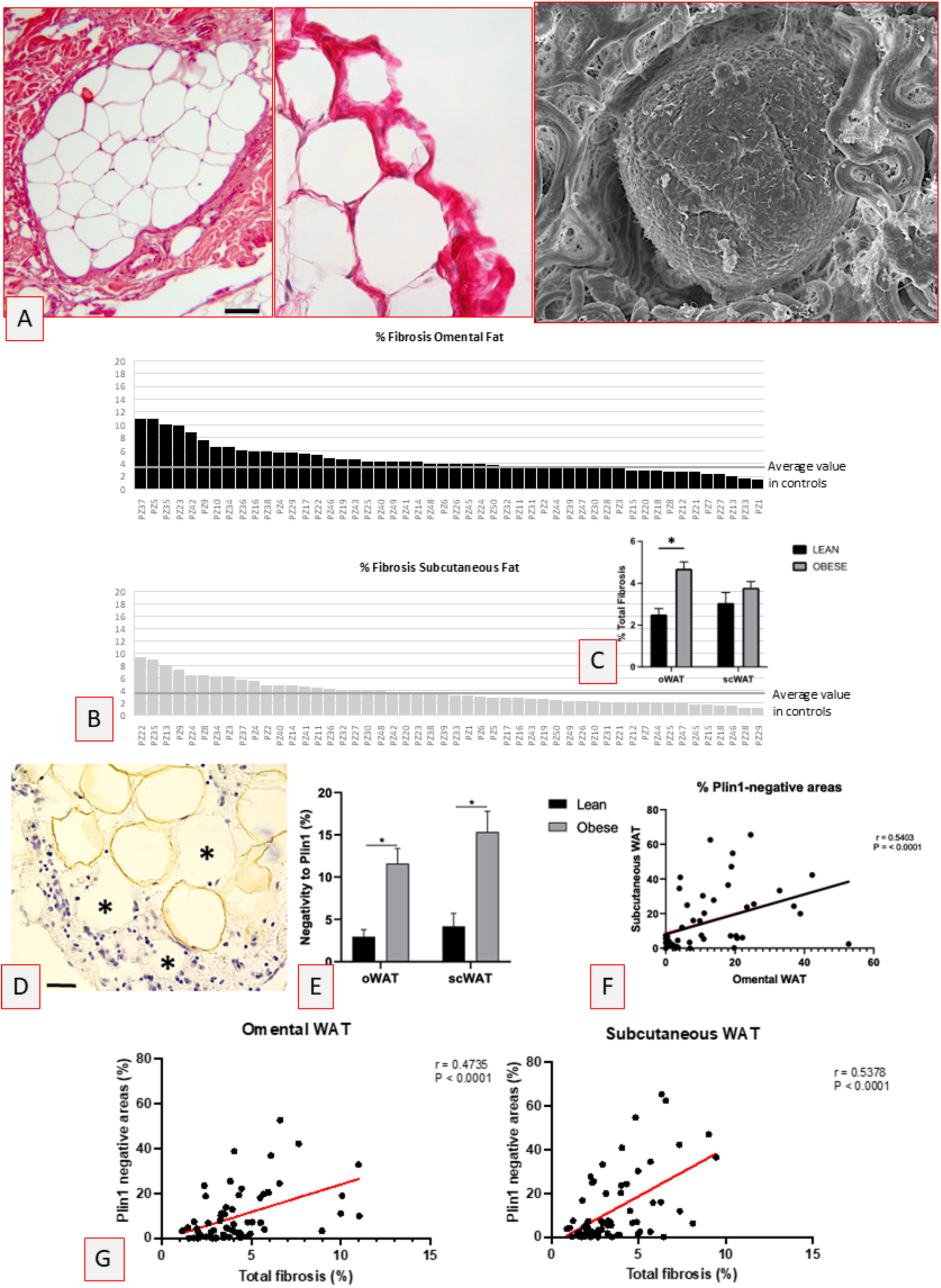
Fibrosis in human fat. A: Light and electron microscopy. Sirius Red staining (left and middle panels): representative images of fibrosis in visceral (omental) fat. In the right panel pericellular fibrosis is shown by high resolution scanning electron microscopy (HRSEM see methods). Bar: 50μm (left), 15 μm (middle) and 7μm (right). B: Percentage of parenchymal fat occupied by the Sirius Red-stained fibrotic tissue in each patient with obesity (see methods for details). C: Fibrosis resulted significantly increased only in omental fat when compared with lean controls. D: Perilipin1 (PLIN1) immunohistochemistry. PLIN1-negative adipocytes (*) close to PLIN1 immunoreactive adipocytes are visible in a visceral biopsy from a patient with obesity. Bar: 40 μm E: The percentage of PLIN1- negative parenchymal areas resulted higher in both subcutaneous and omental fat from patients with obesity. F: Correlation between PLIN1-negative parenchymal areas of visceral and subcutaneous fat in each patient. G: Correlations between percentage of PLIN1-negative parenchymal areas and total fibrosis in subcutaneous and omental fat.

**Fig 2.**
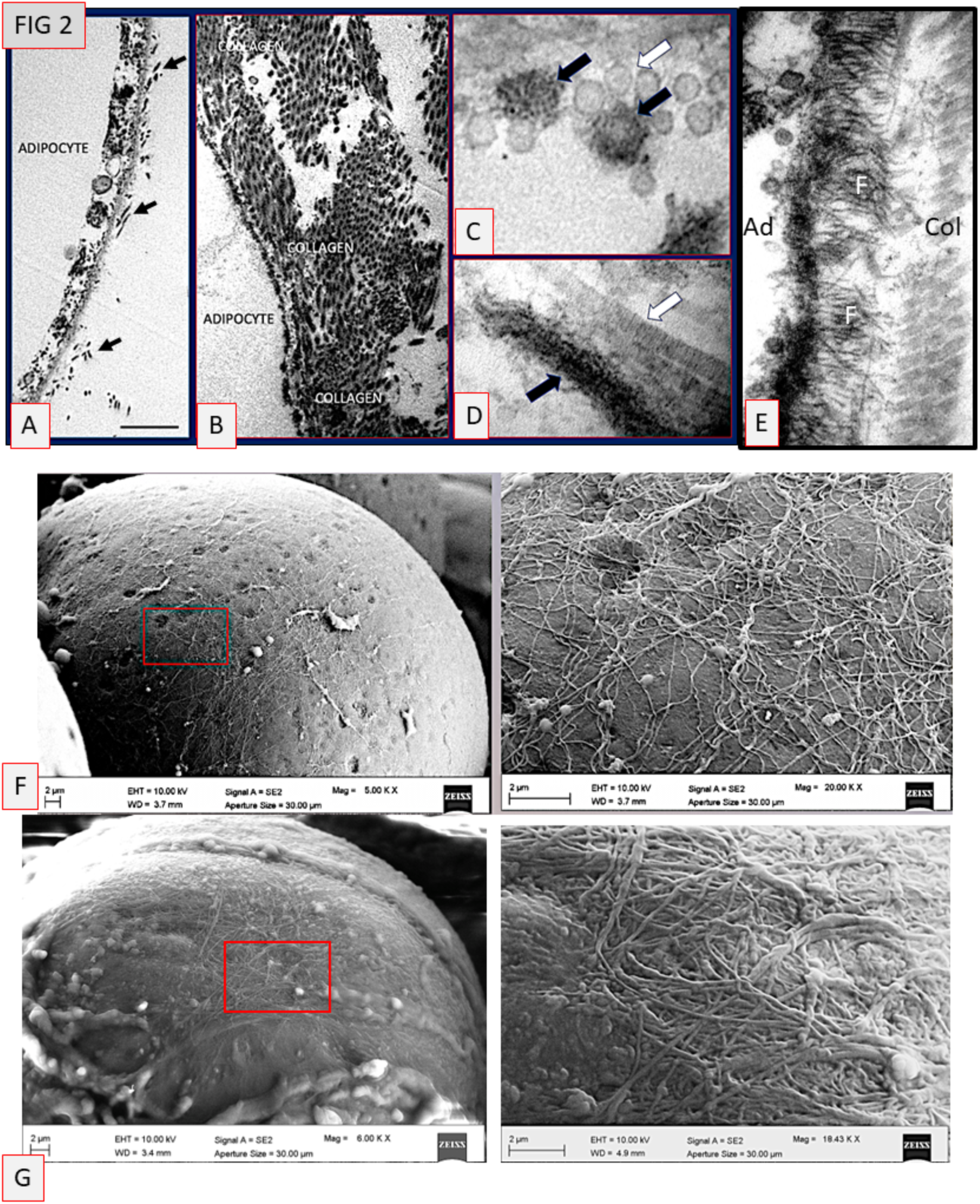
Transmission (TEM) and scanning (high-resolution) (HRSM) electron microscopy of human visceral (omental) fat. A: TEM: lean control, note few sparse collagen fibrils in close contact with the external surface of the adipocyte (arrows). B: TEM: representative biopsy from a patient with obesity, note the abundant collagen fibrils tightly connected with the adipocyte. C: high magnification of transverse sectioned collagen fibrils shown in B, note the enlarged and dense fibrils (black arrows) and compare with normal fibrils (white arrow). D: longitudinal section of collagen fibrils showing an altered fibril (black arrow) and a normal fibril (white arrow) morphology. E: Representative image of hypertrophic adipocyte (Ad) from obese fat with microfibrils (F: possibly tropocollagen) apparently sprouting directly from the cell surface. Col: collagen fibrils. Bar: 1 μm in A, 1.8 μm in B, 115 nm in C, 230 nm in D and 500 nm in E. F and G: Representative images of High-Resolution Scanning Microscopy (HRSEM) of omental fat from a lean patient (F) and a patient with obesity (G) with fibrosis >5%. Right panels are enlargements of squared areas in the left panels. Note the density and thickness of fibrils in the network enwrapping adipocytes in the fat from the patient with obesity. Bars as indicated

The distribution of fibrosis among patients was quite heterogeneous (Fig 1B).

When compared to data from corresponding fat tissue from lean patients, only omental fat fibrosis showed a significant increase (Fig 1C).

Correlations with lipidic metabolic markers (and with a tendency for glycemia) further confirmed the important clinical significance of visceral fibrosis, in agreement with previous studies^3–5,17,35,36^ (TAB IV).

**Table IV.** Correlation analysis between adipose tissue fibrosis and clinical parameters.

|  | <u>Fibrosis in oWAT</u> |  | <u>Fibrosis in scWAT</u> |  |
| --- | --- | --- | --- | --- |
|  | R | P | R | P |
| Age | -0.16 | 0.25 (ns) | 0.2 | 0.16 (ns) |
| BMI | -0.12 | 0.39 (ns) | -0.03 | 0.83 (ns) |
| Adipocyte size ($\mu\text{m}^2$) | -0.33 | 0.022 | -0.35 | 0.01 |
| Triglycerides | -0.37 | 0.02 | -0.18 | 0.24 (ns) |
| HDL cholesterol | 0.36 | 0.02 | 0.11 | 0.48 (ns) |
| LDL cholesterol | 0.35 | 0.03 | -0.09 | 0.55 (ns) |
| Glycemia | -0.27 | 0.06 (ns) | -0.04 | 0.77 (ns) |
| HbA1c | -0.16 | 0.38 (ns) | 0.04 | 0.84 (ns) |

oWAT: visceral (omental) fat
scWAT: subcutaneous (abdominal) fat

### A large proportion of hypertrophic adipocytes are PLIN1 - negative

Hypoxia due to adipocyte hypertrophy is thought to contribute to fibrosis^37,38^. However, the anatomical arrangement of fibrous bundles encasing groups or individual adipocytes could also lead to a form of “self-choking” induced hypoxia (Fig 1A).

Hypoxia can cause adipocyte death, and dead adipocytes surrounded by pericellular fibrosis have been described^36^, though not previously quantified.

Since PLIN1 is a marker of adipocyte vitality^25,39,40^, we measured the percentage of tissue occupied by PLIN1-negative adipocytes.

Surprisingly, an impressive amount of subcutaneous (about 15%) and visceral fat (about 12%) parenchyma from obese patients consisted of PLIN1-negative (dead or distressed) adipocytes compared to only 4% (subcutaneous) and 3% (visceral) in controls (Fig 1D and E). A positive correlation between subcutaneous and visceral PLIN1-negative adipocytes in individual patients was observed, suggesting a general individual phenomenon (Fig 1F). The percentage of PLIN1-negative adipocytes in lean patients matched published data on physiologic adipocyte turnover in humans^41^.

We further noted that many PLIN1-negative adipocytes were located near fibrotic areas. To explore this relationship, we analyzed the correlation between fibrosis and PLIN1-negative adipocytes areas. A strong positive correlation was found (Fig 1G), which remained significant even after stratifying the population by gender (Suppl Fig 2).

These findings suggest that adipocyte stress or death could play a role in the development of fibrosis in patients with obesity.

### Obese adipocytes could play a role in fibrosis

In a previous work on adipose tissue from obese mice, we observed electron microscope images suggesting that adipocytes directly produce collagen fibrils^42^. To further investigate this hypothesis, we analyzed electron microscopy images of visceral fat from patients with obesity and fibrosis >5% (n = 9) and compared them with those with fibrosis <5% (n = 9) and visceral fat biopsies from lean patients (n = 6).

Both TEM and HRSEM images strongly suggested collagen fibril production by adipocytes, with no clear difference between the two groups (>5% or <5%). In lean subjects, TEM showed few collagen fibrils tightly associated with the basal membrane of adipocytes (Fig 2A). However, in obese patients, a layer of collagen fibrils of variable thickness and in tight connection with the basal membrane of each single adipocyte was often present (Fig 2B). Moreover, among the collagen fibrils with normal anatomy, i.e. mainly type I and III, we found, in visceral fat, a variable number of fibrils with visually evident alterations mainly in patients with fibrosis >5%. Their diameter was twice that of normal type I and III collagen fibrils (100-150 vs 30-70 nm) and their ultrastructure appeared frayed (Fig 2C, D). These data are in line with those found in lean and obese mice^42^, including the frequent appearance of direct fibrillar sprouting from hypertrophic adipocytes. Of note, the fibrillar sprouting was formed by nanofibrils of about 15 nm of diameter, thus about ¼ of that of fibrillar collagen, suggesting their tropocollagen nature (Fig 2E)^43^.

HRSEM further confirmed the presence of fibrils on the adipocyte surface and revealed their organization as a network enveloping the surface of the whole lean adipocyte, both in subcutaneous and visceral fat, thus suggesting that this collagen fibril network own to the normal anatomy of adipocytes (Fig 2F). In line with the TEM data described above, this fibrillar network proved thicker and made up of larger fibrils (with larger ones around 150-200 nm) in hypertrophic adipocytes in individuals with obesity (Fig 2G).

### Human adipocytes express collagen genes in vitro

All together these data suggest the idea that at least part of the collagen fibrils responsible for fat parenchymal fibrosis could be due to direct production by adipocytes. To confirm this hypothesis, we analyzed gene expression in a well-known human adipose cell model: hMADS^26^ and, considering the widely accepted notion that collagen fibrils are produced by fibroblasts, we compared data obtained by a human dermal fibroblast cell line^28^.

The results showed that differentiated hMADS (corresponding to mature adipocytes) express similar levels of gene expression of *COL1A1*, but more *COL3A1* and *COL6A3* than fibroblasts (NhDF) (Fig 3A).

**Fig 3.**
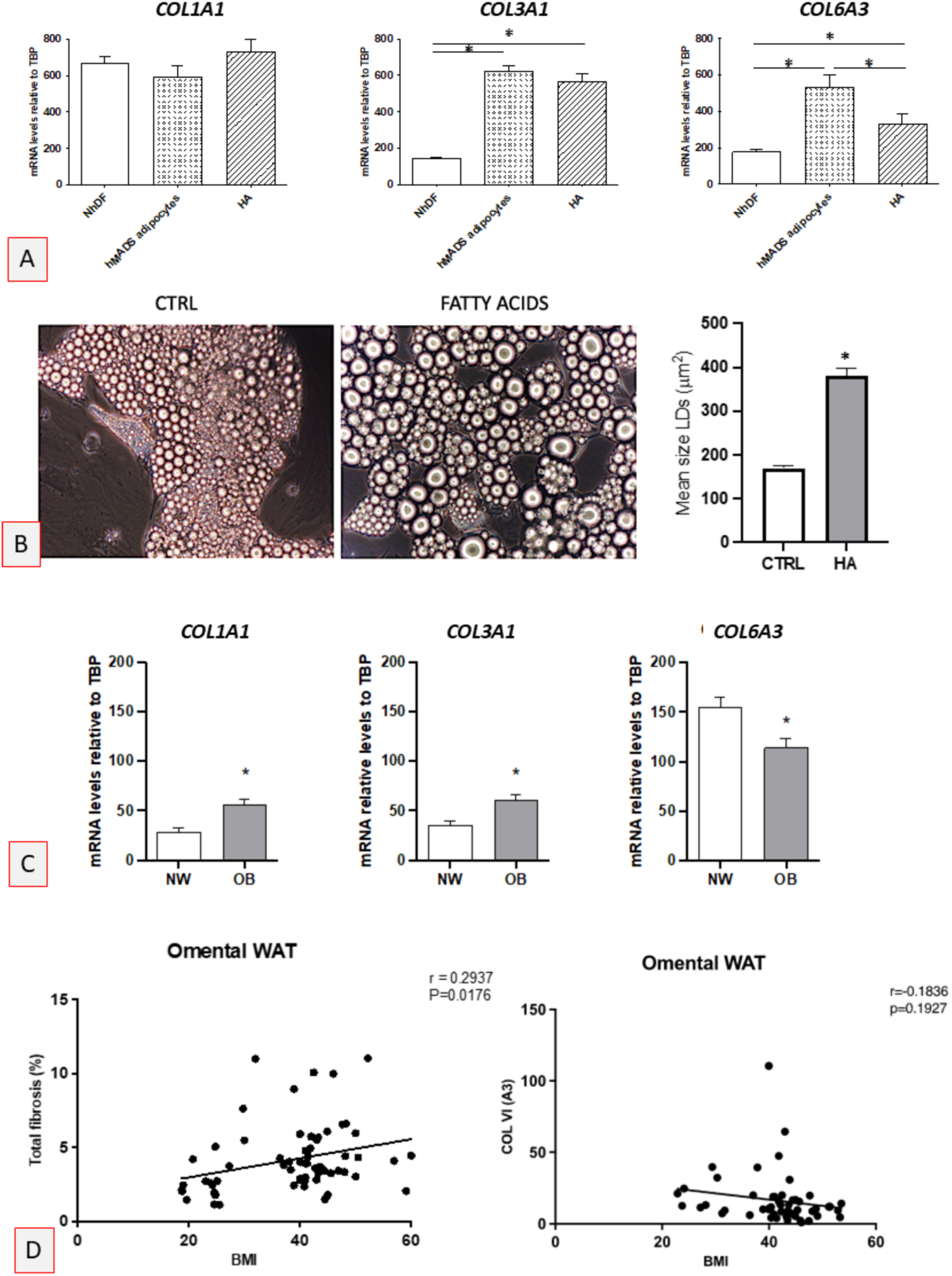
Collagen genes expression. A: In vitro gene expression of COL1A1, COL3A1 and COL6A3 in a human adipocyte cell line (hMADS) and in human dermal fibroblasts (NhDF). Mature adipocytes express the same amount of COL1A1 but more COL3A1 and COL6A3 than that expressed by NhDF. Hypertrophic adipocytes (HA) vs mature adipocytes express more COL1A1 (ns) and less (p<0.05) COL6A3. B: Light microscopy of differentiated hMADS. The addition of fatty acids to the medium induced hypertrophy of lipid droplets (LDs), thus mimicking the phenotype of hypertrophic obese adipocytes (HA) (left and middle panels). Mean size of LDs resulted significantly increased in differentiated hypertrophic hMADS (right panel, p<0.05). C: Increased gene expression of types I (COL1A1) and III (COL3A1) collagens (p<0.05) and reduced gene expression of COL6A3 (p<0.05) in visceral adipose tissue from patients affected by obesity (OB n 50) and lean controls (NW n 15). D: Omental (visceral) fat: Positive correlation between BMI and fibrosis (Left). Negative correlation between BMI and COL6A3 (right). Of note: data from lean patients are included.

We then sought to develop an adipocyte condition like that of adipocytes in obese fat. We added fatty acids to the culture medium in order to obtain hypertrophic adipocytes^27^ (HA). Mean size of lipid droplets was measured (Fig 3B) and the collagen gene expression in these cells was verified (Fig 3A). Interestingly, all collagen types were expressed in hypertrophic-like adipocytes, with a clear tendency for increased expression of *COL3A1* when compared to fibroblasts. Of note, *COL6A3* expression in hypertrophic adipocytes decreases when compared with normal mature adipocytes, in line with data obtained *in vivo* from fat biopsies (see below). All together these data further suggest that obese adipocytes produce collagen fibrils and could contribute to fibrosis.

### Increased intra-abdominal pressure can be the stimulus for visceral fat fibrosis

Next, we asked which stimulus could be responsible for collagen production in hypertrophic adipocytes.

Adipocytes are mechanosensitive and mechanoresponsive^44^. We turned to the role of pressure because adipose tissue under strong pressure, such as that of plantar calcaneal fat, has a peculiar anatomy including abundant pericellular collagen^45^, confirmed also by our original data (Suppl Fig 3). Thus, we investigated if abdominal pressure, acting mainly on visceral fat, is increased in patients with obesity. In line with old published data^46^, we found increased intra-abdominal pressure in subjects suffering from obesity with a positive correlation with abdominal circumference (visceral obesity) (Suppl Fig 4).

Thus, increased pressure on adipocytes could trigger the molecular mechanism responsible for increased collagen production and subsequent fibrosis.

### Obese fat is inflamed

The death of adipocytes^25^ can cause fat inflammation, especially in visceral fat^34,40^. Thus, we checked for inflammatory cells. CD68 immunohistochemistry showed that the vast majority of inflammatory cells in both visceral and subcutaneous obese fat were macrophages, known to trigger fibrosis^47^. Both tissues proved to be inflamed with a positive correlation for single patients and with more macrophages in visceral than in subcutaneous fat (Suppl Fig 5A).

Considering that >90% of macrophages in obese mice are organized to form crown-like structures (CLS)^25^, we sought to determine the localization of macrophages in inflamed tissues. We found that only a minority of macrophages was localized in CLS. Most macrophages were localized among apparently normal adipocytes in parenchyma, although a conspicuous amount of them were localized in fibrotic areas, or within or near vascular structures (Suppl Fig 5 B). Dead adipocytes can cause inflammation^25,34,40^, but we have not found a correlation between PLIN1-negative (dead or suffering adipocytes) and the number of macrophages (data not shown). However, we did find a positive correlation with fibrosis, in line with the well-known pro-fibrotic activity of macrophages^47^ (Suppl Fig 5 C).

These data suggest that inflammation plays a key role in fibrosis, in line with data from other studies^17,48^.

### Molecular mechanisms: role of Collagen VI

While data on increased Col I and III in obese human fat are largely confirmed^35,36^, data on Col VI remains controversial^10–13^.

Gene expression analyses in our patients revealed that obese visceral fat showed a decrease in *COL6A3* expression, while *COL1A1* and *COL3A1* levels were elevated (Fig 3C). Likewise, we found an evident reciprocal relationship between BMI and fibrosis (positive) and BMI and *COL6A3* gene expression (negative) (FIG 3D).

We therefore investigated whether silencing *COL6A3* in hMADS affected the gene expression of *COL1A1* and *COL3A1*. Our results showed an increased expression of both types of collagen genes after 7 days in culture in cells with about 50% of *COL6A3* gene silenced. Specifically, *COL1A1* increased by 75% and *COL3A1* by approximately 30% (Fig 4A).

**Fig 4.**
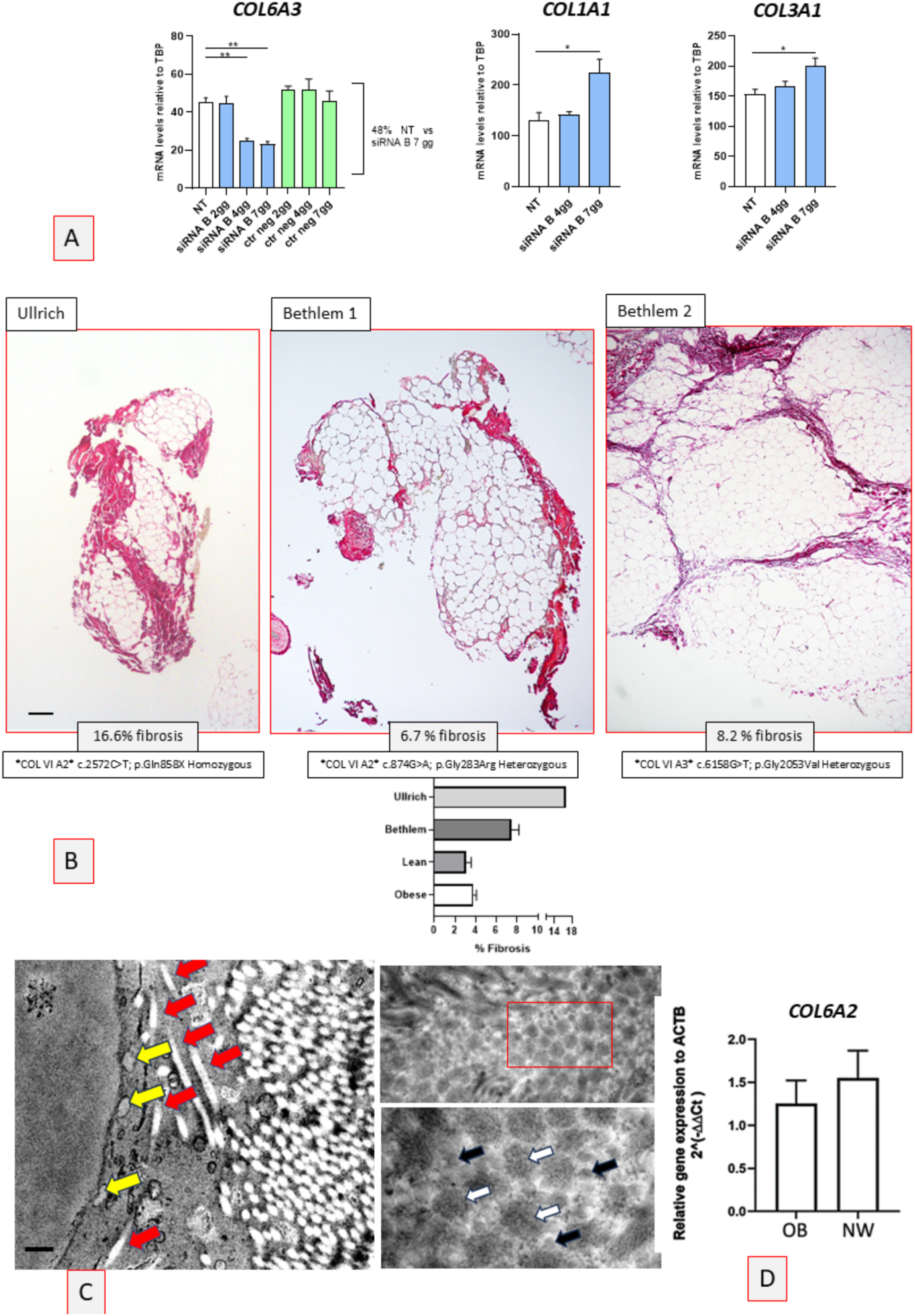
Gene silencing in human adipocytes (hMADS) in vitro and human Col VI congenital gene mutations in vivo seem to induce fibrillar collagen production by adipocytes. A: hMADS with silenced 48% of COL6A3 gene show increased production of COL1A1 and COL3A1 after 7 days. B: Representative light microscopy with Sirius Red staining of fat biopsies from Ullrich case (left panel) and Bethlem cases (middle and right panels). Details of Col VI genes mutations are indicated. Note the abundant lobular and pericellular fibrosis evident in all cases. Bar: 80μm (right panel), 45μm (middle panel) and 50μm (right panel). A summary of the fibrosis percentage found in subcutaneous fat from studied patients is shown in a graph. C: Representative transmission electron microscopy (TEM) image showing an impressive amount of fibrillar collagen in tight contact with a mature adipocyte (Ullrich case, see also Suppl Fig 6). Note the location of some mature collagen fibrils (red arrows) into the cytoplasm (left panel). Rough endoplasmic reticulum lumen is dilated (yellow arrows). Right panels showing transverse sections of collagen fibrils. Upper right panel: some collagen fibrils resulted enlarged. Lower right panel: enlargement of squared area in right upper panel. Compare enlarged (white arrows, some indicated) with normal fibrils (black arrows, some indicated) Bar: 200 nm in left panel, 450 nm in upper right panel and 230 nm in right lower panel. D: COL6A2 tended to be less expressed in visceral fat of patients suffering for obesity (OB n 25) than in lean patients (NW n 12).

### Patients with mutations in COL6A2 and COL6A3 genes

We had the opportunity to analyze the adipose tissue from subcutaneous fat of three patients affected by COLVI-RMs, one with a *COL6A2* homozygous mutation, with UCMD, and two patients with BM caused by heterozygous mutations in *COL6A2* (BM1) and *COL6A3* (BM2).

All three patients were lean by BMI (clinical data in TAB III), and the size (area) of their adipocytes was in the normal range considering also the patient’s age^49^: about 1,900μm^2^ Ullrich (11 years old patient) and about 4,590μm^2^ (patient 1, 40 years old) and 3,870 (patient 3, 41 years old) with average of 4,200 +/- 360μm^2^ vs. about 4,700 μm^2^ in lean control patients (difference not significant).

In all patients, we found an impressive amount of fibrosis, quantified by Sirius Red-stained collagen fibers in parenchymal fat using the same method used to quantify fibrosis in patients with obesity (see Methods for details). Fibrosis was more evident in the Ullrich patient (16.6% of parenchymal fat) than in the Bethlem patients (6.7% patient 1 and 8.2% patient 3), thus all well above that found in control lean patients (2.5%) (Fig 4B).

PLIN1 immunoreactivity showed 5% of negative adipocytes in the Ullrich case and about 7% (patient 1) and 7.9% (patient 2) in Bethlem cases (data not shown). Thus, despite the high level of fibrosis, PLIN1-negative adipocytes were less than the average in obese fat (about 15%); it is possible that the size of adipocytes in these patients (see above) could account for the difference. CD68 immunostaining and TEM excluded significant signs of inflammation in all three patients (data not shown).

TEM showed in all cases a tight relationship between adipocytes and collagen fibrils that, together with hypertrophy of organelles for protein synthesis and secretion, strongly suggested a direct role of these cells in collagen production (Fig 4C left panel). Notably, in the Ullrich patient, altered, enlarged fibrils, resembling those found in obese fat fibrosis, were observed (Fig 4C right panels and Suppl Fig 6), supporting the hypothesis that Col VI plays an important role in the stabilization of collagen fibrils, as suggested by other studies^50^.

Together, these data highlight the significant role of Col VI in adipocyte-driven fibrosis. Furthermore, considering that in the patients with obesity of this case series we found a reduction (about 30%) in the gene expression of *COL6A3* (corresponding to that found in patient Bethlem 3) we checked also for *COL6A2* (corresponding to that found in Ullrich patient and Bethlem 1) and found about 15% of reduced expression (Fig 4D).

### Role of CD38

Recent studies have highlighted the significantly higher expression of CD38 in the skin of subjects with systemic sclerosis compared to healthy subjects^18^, suggesting that the CD38 ectoenzyme plays a key role in the onset of tissue fibrosis. We therefore aimed to investigate whether CD38 is expressed in the tissues of patients with obesity.

Immunohistochemical analysis of omental (n = 8) and subcutaneous (n = 2) adipose tissue from obese patients, as well as omental (n = 3) and subcutaneous (n = 3) adipose tissue from lean patients, revealed positivity in some sporadic adipocytes and macrophages in lean subjects, both in subcutaneous and visceral fat. The positive adipocytes exhibited a clear granular-like morphology, suggesting the presence of the intracytoplasmic type of CD38^51^. Interestingly, an area of fibrosis containing several fibroblast-like cells (widely considered responsible for fibrosis) in a subcutaneous biopsy was negative for CD38 (Fig 5A left panels).

**Fig 5.**
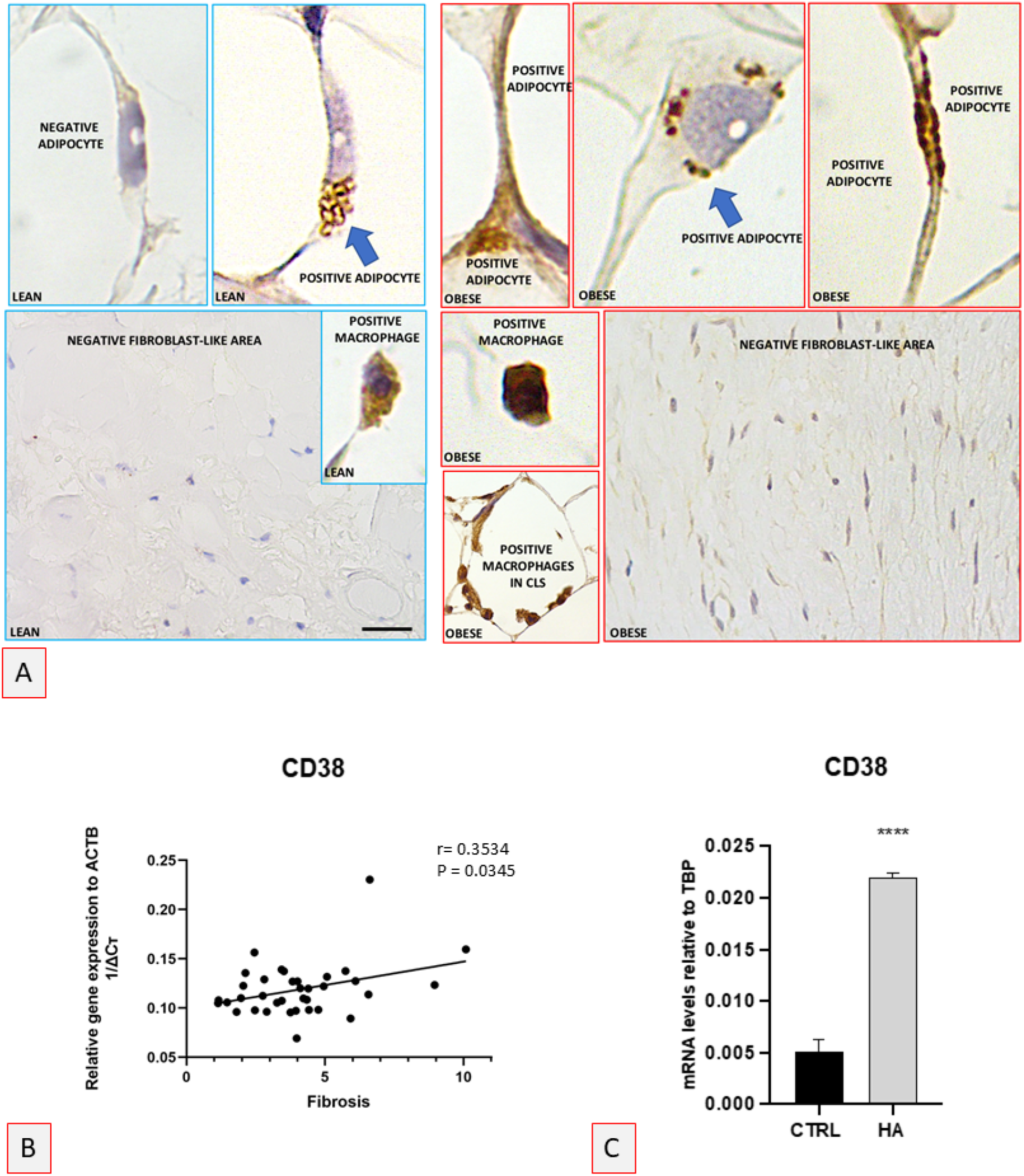
Immunohistochemistry with CD38 antibodies. A: Light microscopy. Representative images from lean (LEAN) or affected by obesity (OBESE) visceral fat. Most of the lean adipocytes were negative. Rare positive adipocytes showed a granular-like positivity (arrow). Weakly positive macrophages were also found in lean fat. Fibrous tissue resulted negative in both lean and obese fat. Intense granular-like positivity in adipocytes was found in fat from patients suffering for obesity. Macrophages resulted intensely positive both when located in interstitial parenchymal tissue as well as when forming CLS in obese fat. Bar: 3μm in all upper panels; 30 µm in lower left and right panels; 10μm in upper middle panels (macrophages), 20 μm in lower middle panel (CLS). B: a positive correlation was found between fibrosis and CD38 gene expression. C: CD38 gene expression in the human adipocyte cell line hMADS in standard condition and after the addition of fatty acids in the culture medium to obtain hypertrophic adipocytes. (See also Fig 3).

In the adipose tissues of subjects affected by obesity, both adipocytes and macrophages in both subcutaneous and visceral fat were frequently intensely positive for CD38. In particular, the immunoreactivity in obese adipocytes was of the granular type. The areas of fibrosis were negative also in these patients (Fig 5A right panels). A positive correlation between *CD38* gene expression and fibrosis was detected (Fig 5B).

Taken together, these data support the overexpression of *CD38* by both macrophages and adipocytes in the adipose tissue of individuals affected by obesity. We therefore investigated whether the human adipose cell line hMADS also expresses *CD38* and whether its hypertrophy was accompanied by increased expression. The data showed that hMADS cells express *CD38*, and its expression increases in the hypertrophic adipocyte model (HA) (Fig 5 C).

Thus, in addition to the role for Col VI, a potential role for CD38 in human fat fibrosis could also be considered. Using mature hMADS, we examined whether a functional relationship between *CD38* and *COL6A2 and A3* exists. Data showed that Col VI genes expression did not change following administration of a *CD38* stimulator at various doses (Suppl. Fig 7).

### Other players in fat fibrosis

Data from other papers suggest that the cells responsible for fibrosis in obese human fat are myofibroblasts and adipocyte precursors (see ^17^ for a recent review). Exploring whether a correlation between our data on fibrosis and markers of these cell types exists, we found a positive correlation with the marker of myofibroblasts αSMA (*ACTA2*), but not for activin A (*INHBA*) (Fig 6A and B). Our findings suggest that fibrotic changes in obese adipose tissue are closely linked to increased expression of myofibroblast markers, particularly αSMA, but not necessarily to other factors such as activin A. This is consistent with the notion that myofibroblasts (characterized by αSMA) are central drivers of pathological fibrosis in adipose tissue, as they actively secrete and remodel extracellular matrix (ECM) components, contributing to the stiffening of the adipose microenvironment ^52,53^

**Fig 6.**
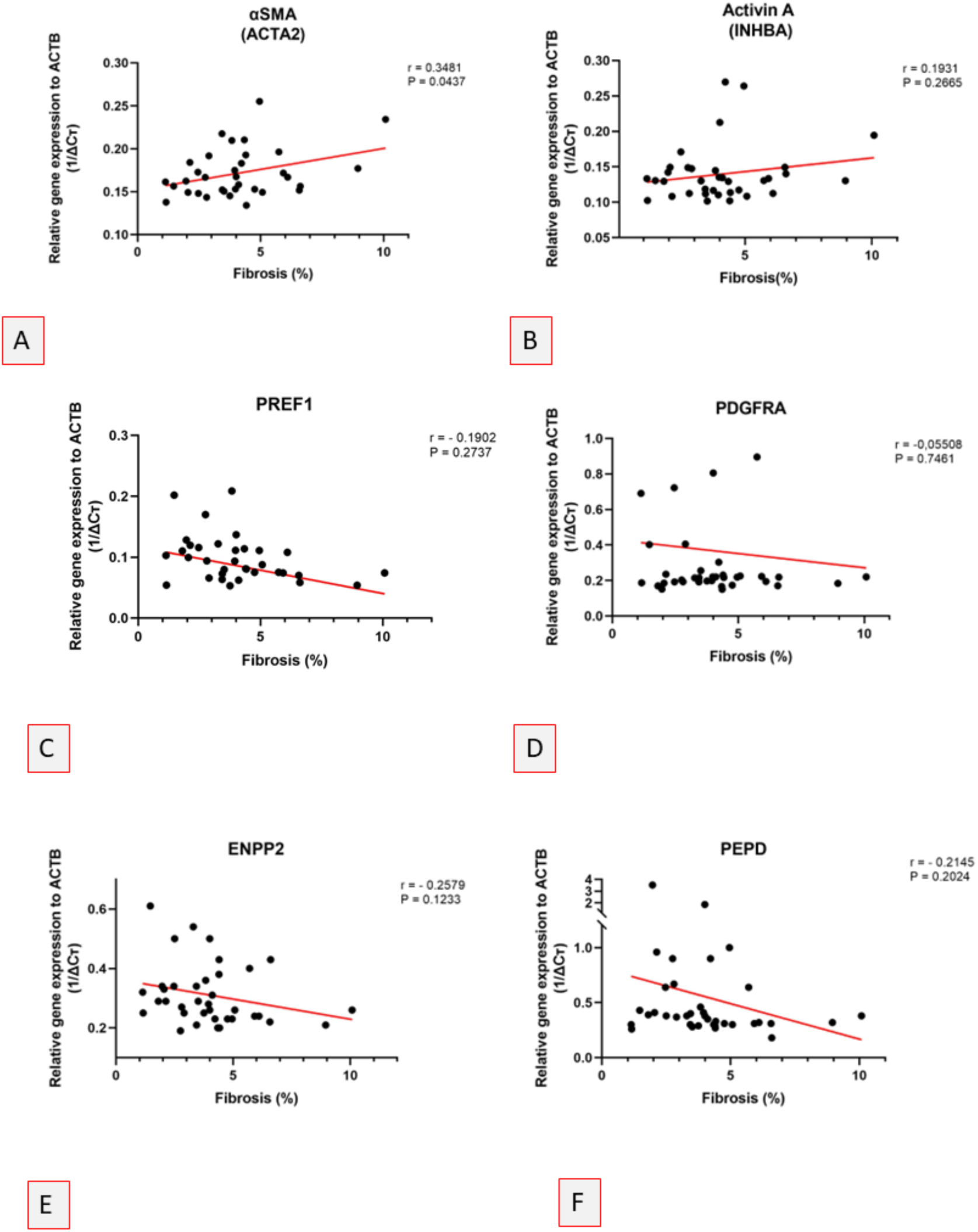
Gene expression in visceral (omental) fat from patients with obesity of our case series. Correlations suggesting a role for myofibroblasts (αSMA with a positive tendency for Activin A) and peptidase D (PEPD)(F) in the pathogenesis of visceral fat fibrosis were found (A, B and F). Data did not support a role for preadipocytes (PDGFRA and PREF 1) and Autotaxin (C, D and E). (OB n 25, lean n 11)

A negative correlation was found with the preadipocyte marker *PREF1* and a negative trend for *PDGRα* (Fig 6C and D). The negative correlation observed with *PREF-1* (preadipocyte factor) and the negative trend with *PDGFRα* suggest that fibrosis may damage adipogenesis by reducing the pool of functional preadipocytes. A fibrotic microenvironment, enriched in collagen and other ECM proteins, can impair adipocyte differentiation, thus reflecting an inverse relationship between fibrotic burden and the capacity of adipose progenitors to undergo healthy adipogenesis^54^. Autotaxin (*ATX*), also known as ecto-nucleotide pyrophosphatase/phosphodiesterase (*ENPP2*) encoded by the *ENPP2* gene, has been suggested to play a role^55^, but we found a tendency for a negative correlation in our study (Fig 6E). The subtle negative correlation with *ATX* may reflect the complex context-dependent role of lysophosphatidic acid (*LPA*) signaling in tissue remodeling. Moreover, we examined the correlation between extracellular enzyme peptidase D (*PEPD*) and fibrosis and found a negative correlation tendency in the visceral fat of our case series (FIG 6F). This supports the idea that PEPD, responsible for collagen degradation and protection against a fibrotic microenvironment, plays a role in fibrosis in line with data from other studies^22^. Reduced expression or activity of PEPD may impair collagen remodeling, thereby facilitating the progression of fibrotic processes.

## Discussion

Our data, obtained through both transmission and high-resolution scanning electron microscopy, indicate, for the first time, that the normal anatomy of human adipocytes includes a fibrillar network surrounding each cell, similar to that previously described in mice^42^.

The function of this network is probably linked to the expansibility needs of adipocytes, providing elastic mechanical support that counteracts the cell expansion strength^16,56^. On the other end this implies a physiologic need of fibrillar collagen production by adipocytes.

Our data support that fibrillar collagen (I and III) is increased in human obese visceral fat, in line with data from other studies^35,36^, but also suggest that it is directly produced also by adipocytes.

The cell origin of this collagen is widely considered to be fibroblasts or fibroblast-like cells (myofibroblasts)^17^. However, our electron microscopy data suggest that obese adipocytes may also serve as a source. Compared to human dermal fibroblasts, mature human adipocytes express *in vitro* similar levels of *COL1A1*, but more than three times the level of *COL3A1*, with a tendency for increased expression under hypertrophic-like conditions. These findings, together with the electron microscopy results, suggest a direct fibrillar production by obese adipocytes and support the idea that adipocytes themselves may be a significant cellular source of fat fibrosis. This hypothesis is in line with recent proteomics data showing that isolated visceral adipocytes from mice, under high fat diet, adopt a fibroblast-like phenotype^57^.

Of note, the presence of intracytoplasmic collagen fibrils we found in the adipocytes of Ullrich patient, with the highest levels of fat fibrosis (Fig 4C and Suppl Fig 6) further support the direct collagen production by adipocytes. The presence of intracytoplasmic fibrils in cells with high levels of collagen production is well-documented in a case of desmoid fibromatosis, in which several fibroblasts showed intracytoplasmic collagen fibrils^58^.

Data on Col VI abundance in human obese fat are controversial^10–13^. Our data show a reduced expression of *COL6A2 and COL6A3* mRNA transcripts. In humans, mutations in *COL6A1*, *COL6A2 and COL6A3* primarily result in muscular consequences^50,59^. However, our data show that both UCMD and BM patients, with well-known reduced functionality of Col VI^15^, have clear fibrotic consequences in adipose tissue. Subcutaneous fat in these patients exhibited fibrosis levels that were about 6.5 times higher (Ullrich: homozygote case) and 2.8 times higher (Bethlem: heterozygote cases) than those observed in control biopsies from lean patients. Of note, in our patients with obesity, subcutaneous fat did not show a significant increase in fibrosis compared to lean controls, making the fibrosis observed in the Ullrich and Bethlem patients even more striking.

These findings support the notion that, in presence of altered Col VI activity, adipose tissue produces more fibrillar collagen. This is also in line with our in vitro results showing an increased expression of *COL1A1 and COL3A1* after a 50% silencing of *COL6A3* in mature human adipocytes (present paper). Collagen VI is the most represented type of collagen in adipose tissues together with the fibrillar types I and III^7,17^. Its function is still unknown, but data suggest a key role in fibrillar collagen stability^50^, functioning like a glue that interconnects the fibrillar network. Leptin production increases with adipogenesis^60^, and has been shown that it is able to inhibit Col VI^13^, thus it is plausible that leptin reduces Col VI in order to adjust the strength of the fibrillar network surrounding each adipocyte and therefore allowing the cell expansion.

Data from mice lacking Col VI did not reveal any association with fat fibrosis^10^, but it is also known that the phenotype is milder than in the human disorder, and in three different models of mice lacking Col VI, muscle fibrosis has been described^59^.

Pericellular fibrosis is physiologically present in calcaneal fat, where body weight pressure plays a role in the need for elastic properties in this anatomical component of the human foot. In line with this, the intra-abdominal pressure in patients with obesity is increased (present paper and ^46^), with a positive correlation to abdominal circumference (i.e. visceral obesity)^61^. Thus, increased intra-abdominal pressure may inhibit Col VI, leading to reduced stability of the fibrillar network and a subsequent hyperproduction of Col I and III, contributing to fat fibrosis. A sign of reduced stability in the fibrillar network is also reflected in the altered morphology of collagen fibrils, which were enlarged and exhibited a frayed ultrastructure. Notably, these alterations were observed more frequently in fat biopsies from patients with higher levels of fibrosis and in the patient with Ullrich syndrome.

Adipocyte death and fibrosis are key drivers of inflammation^25,40^. Our previous works ^40^, supported by other studies^39^, suggested that the absence of perilipin 1 (PLIN1) is a hallmark of adipocyte death, which triggers macrophage inflammation^62^. The present study shows that a large proportion of obese adipocytes (approximately 15%) lack PLIN1, further supporting the idea that dead or distressed adipocytes may be co-responsible for the inflammation observed in obesity. Most inflammatory cells found in obese fat were CD68-positive, thus belonging to the category of macrophages. In fat from obese mice, more than 90% of macrophages are localized to crown-like structures (CLS)^25^; however, in the present study, only a minority of macrophages formed CLS. The majority were parenchymal and were found to correlate with areas of fibrosis, consistent with the pro-fibrotic activity of macrophages^5^ and in line with other studies^63^. This suggests a potential vicious cycle: adipocyte death → inflammation → fibrosis → further adipocyte death.

Recent studies have highlighted the role of the ectoenzyme CD38 as a key inducer of systemic fibrosis^18^, with macrophages playing a key role in the general scheme of pathogenetic events ending in fibrosis^19^. In particular, the overexpression of CD38 is paralleled by a depletion of NAD+, which in turn reduces the functionality of various substrates, including PARPs (polyADP-ribose polymerases) and sirtuins^64,65^. PARPs perform important DNA repair functions, while sirtuins are involved in the normal stability and metabolism of mitochondria. Macrophages produce cytokines which, with autocrine and paracrine stimulation on fibroblasts, determine overexpression of CD38. DNA repair failure and mitochondrial instability promote inflammation and cellular senescence (inflammageing) resulting in fibrosis^66^. Our immunohistochemistry data support a direct role for macrophages in CD38 production. Both lean and obese adipose tissues, when containing fibroblast-like cells, showed CD38 negativity, despite strong immunoreactivity in macrophages and in line with data from other Authors^67,68^. These findings indicate that CD38 plays a role in visceral fat fibrosis, as its immunoreactivity was increased in both obese adipocytes and macrophages (which are themselves increased in number). Moreover, we observed a positive correlation between *CD38* gene expression and fibrosis levels. We also found CD38 expression in mature adipocytes in vitro, with a significant increase under hypertrophic-like experimental conditions. To investigate the relationship between CD38 and Col VI, we tested their interaction in mature adipocytes *in vitro* but found no correlation. This suggests that these two factors likely play independent roles in the pathogenesis of fat fibrosis with CD38 activity primarily attributed to macrophages and Col VI to adipocytes. This is also in line with the striking evidence of high levels of fat fibrosis in patients with mutations in Col VI genes in absence of any evidence of significant macrophages fat infiltration.

Another important consequence of macrophage infiltration into obese visceral fat is the well-known induction of *TGF-β* by TNFα, which is produced by macrophages^69^. TGF-β is mainly produced by cells of the immune system, including macrophages, and this factor, which is a primary factor promoting tissue fibrosis, is highly elevated in obese adipose tissue both in mice and humans^21^.

Hypoxia induces HIFα, which stimulates fibrosis^20^ and treatment with the HIFα- selective inhibitor PX-478 has been shown to alleviate fibrosis, inflammation, and glucose intolerance^35^. Our data confirm higher levels of HIF1α in obese visceral fat (data not shown), suggesting that hypoxia could be due to pericellular and peri lobular fibrosis (self-choking induced hypoxia by adipocytes). The impressive percentage of PLIN1 immuno-negative adipocytes (about 15%) and the strong correlation with fat fibrosis found in this study point to the relevance of this mechanism.

Other factors contributing to fat fibrosis, such as reduced peptidase activity responsible for collagen degradation^22^ and myofibroblast activity^3^ align with our findings of a negative correlation tendency for *PEDP* and a positive correlation with *αSMA* in fat fibrosis. Autotaxin, a secreted enzyme that generates LPA, a bioactive lipid that influences cellular signaling and function upon binding to G protein-coupled receptors, is produced and released by human adipocytes and upregulated in obesity. It has been proposed to play a role in fat fibrosis^70^. However, we did not observe any correlation between its gene expression and fat fibrosis in our case series study, although the high variability among patients could account for this difference.

## Conclusions

Our data provide compelling evidence for the direct involvement of mature adipocytes in the pathogenesis of visceral fat fibrosis in individuals with obesity. We propose that vicious cycles play a significant role in this pathology: fat accumulation induces increased abdominal pressure, this seems to lead to a reduction in Col VI genes expression in adipocytes, which in turn triggers the hyperproduction of fibrillar collagen (Col I and III). This is supported by both our in vivo and in vitro findings, as well as by data from patients with congenital homozygous and heterozygous Col VI genes mutations. The resulting pericellular and lobular fibrosis could trigger cellular and tissue hypoxia, giving rise to a vicious cycle: adipocyte hypertrophy → increased IAP → reduced Col VI → increased Col I and III → self-choking and death of adipocytes → hypoxia and inflammation (macrophage infiltration) → TGF β → CD38 hyperproduction → increased fibrosis (Scheme 1).

These findings highlight novel molecular players in the pathology of obesity-related visceral fat fibrosis, particularly the roles of Col VI in adipocytes and CD38 in inflammatory macrophages. Given that CD38 antibodies are already in clinical use for other pathologies^71,72^, these insights may have therapeutic implications for treating obesity-related fibrosis.

## GRANTS

PRIN 2017 to Sav.C. and A.G, grant number #2017L8Z2, and Foundation of Molecular Medicine and Cellular Therapy Marche Polytechnic University, Ancona, Italy.

Pia.Ce.Ri research development plan to Ser. C., Department of Medical, Surgical Sciences and Advanced Technologies “GF Ingrassia” (DGFI) University of Catania, # 2020-2022 Catania, Italy

National Council for Scientific and Technological Development (CNPq-Brazil) - (200295/2022-5) to F. C. C.

## Supplementary figures

**SFig 1:**
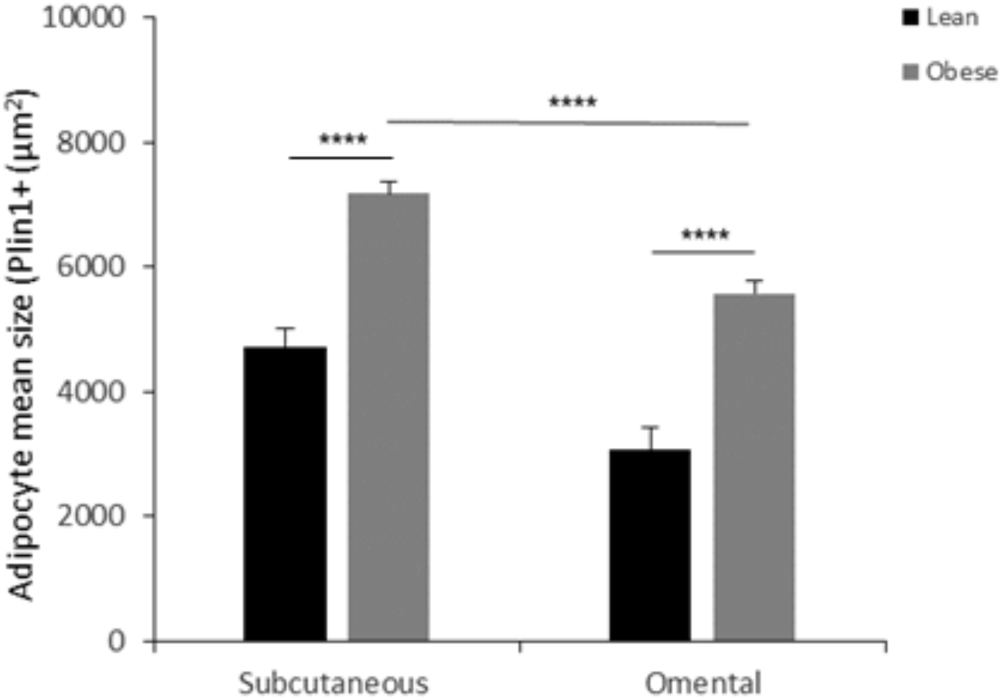
Size of perilipin1 immunoreactive adipocytes in omental and subcutaneous fat of lean and patients with obesity.

**SFig 2:**
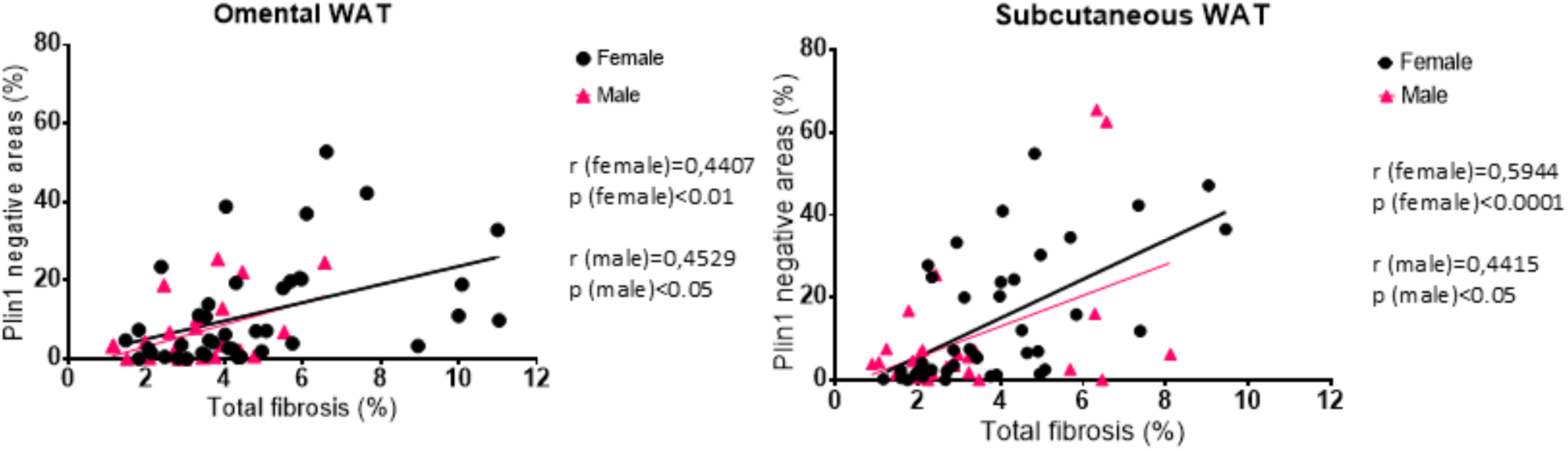
Positive correlation between density of perilipin1 negative adipocytes and percentage of fibrosis in adipose tissue of patients with obesity distinguished by gender.

**SFig 3:**
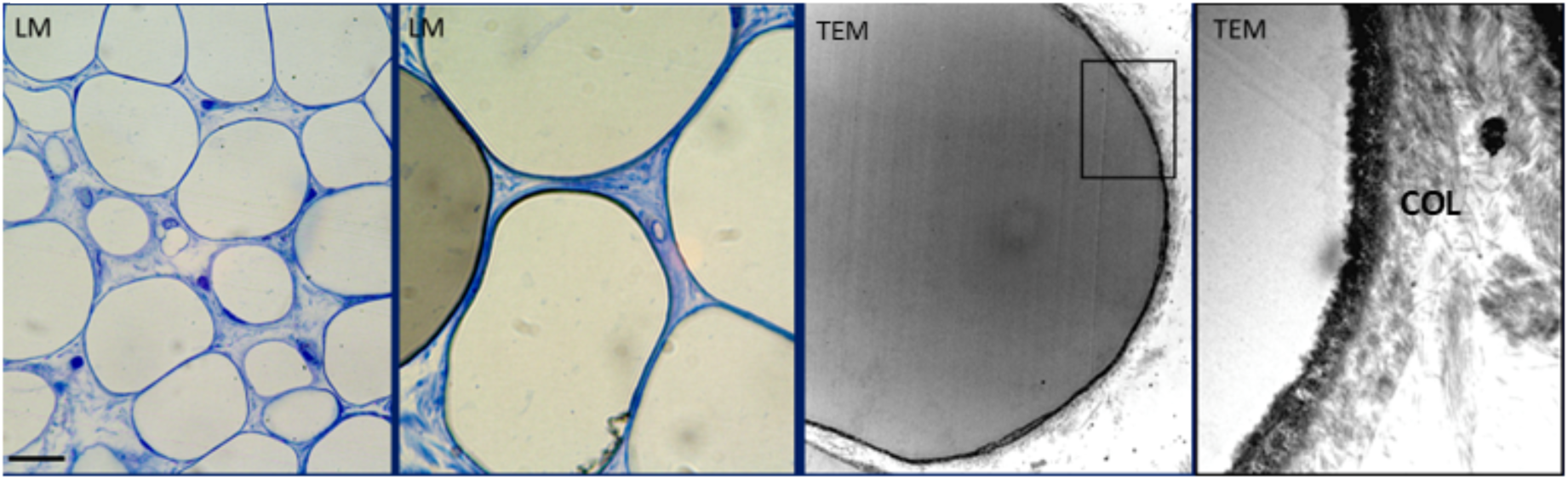
Representative light (LM) and transmission electron microscopy (TEM) images from normal human calcaneal fat. A thick layer of collagen fibrils surrounds each adipocyte: thick blue line in LM (two panels on the left) and Col in the last right panel (enlargement of squared area in the middle right panel). Bar: 10μm in first panel, 4μm in second panel, 2.5μm in third panel and 0.4μm in forth panel.

**SFig 4:**
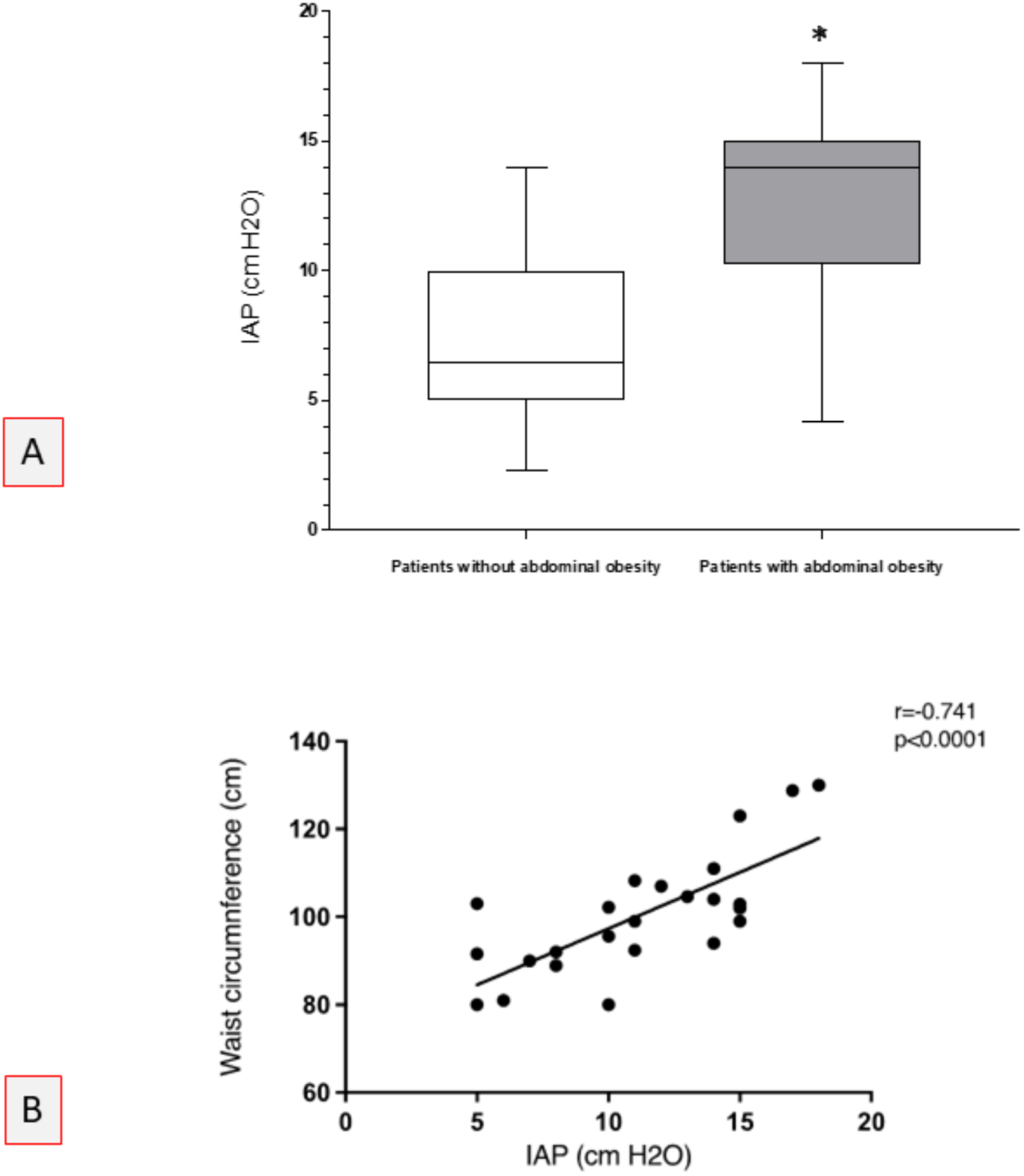
Intra-abdominal pressure (IAP) in humans. A- IAP of patients with obesity resulted increased (p<0.05). B- Correlation between waist circumference and IAP in humans (p<0.001).

**SFig 5:**
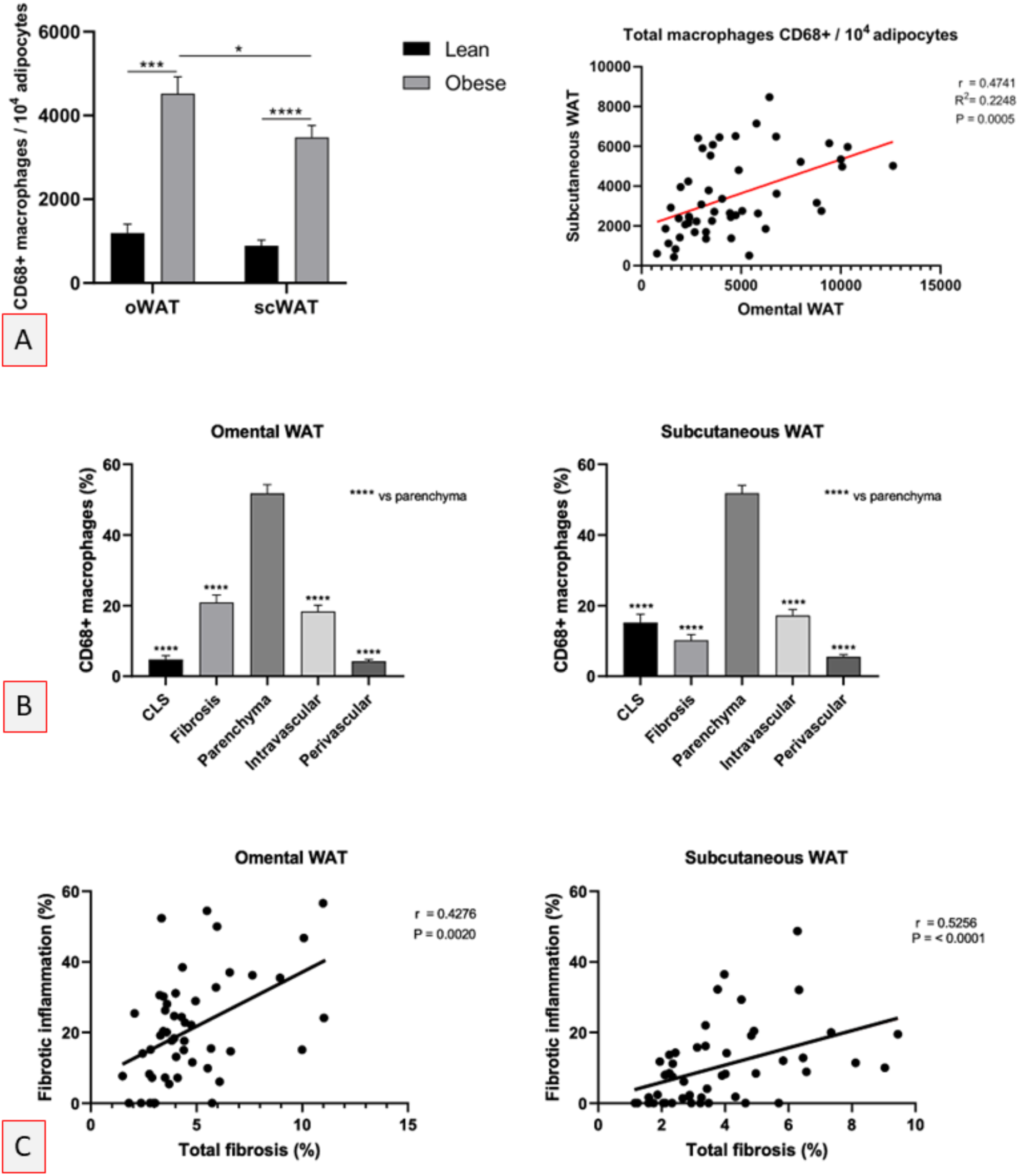
A- Quantitative data showing that CD68 immunoreactive macrophages were more abundant both in subcutaneous and visceral (omental) fat of patients with obesity (upper left panel) with a positive correlation between the two fat compartments per each single patient (upper right panel). B- Distribution of CD68 immunoreactive macrophages in different anatomical sites of adipose tissues in patients with obesity showed that only a minority was localized in crown-like structures (CLS). C- A positive correlation between extension of fibrosis and density of CD68 immunoreactive macrophages in fibrotic areas was found both in visceral and subcutaneous fat.

**SFig 6:**
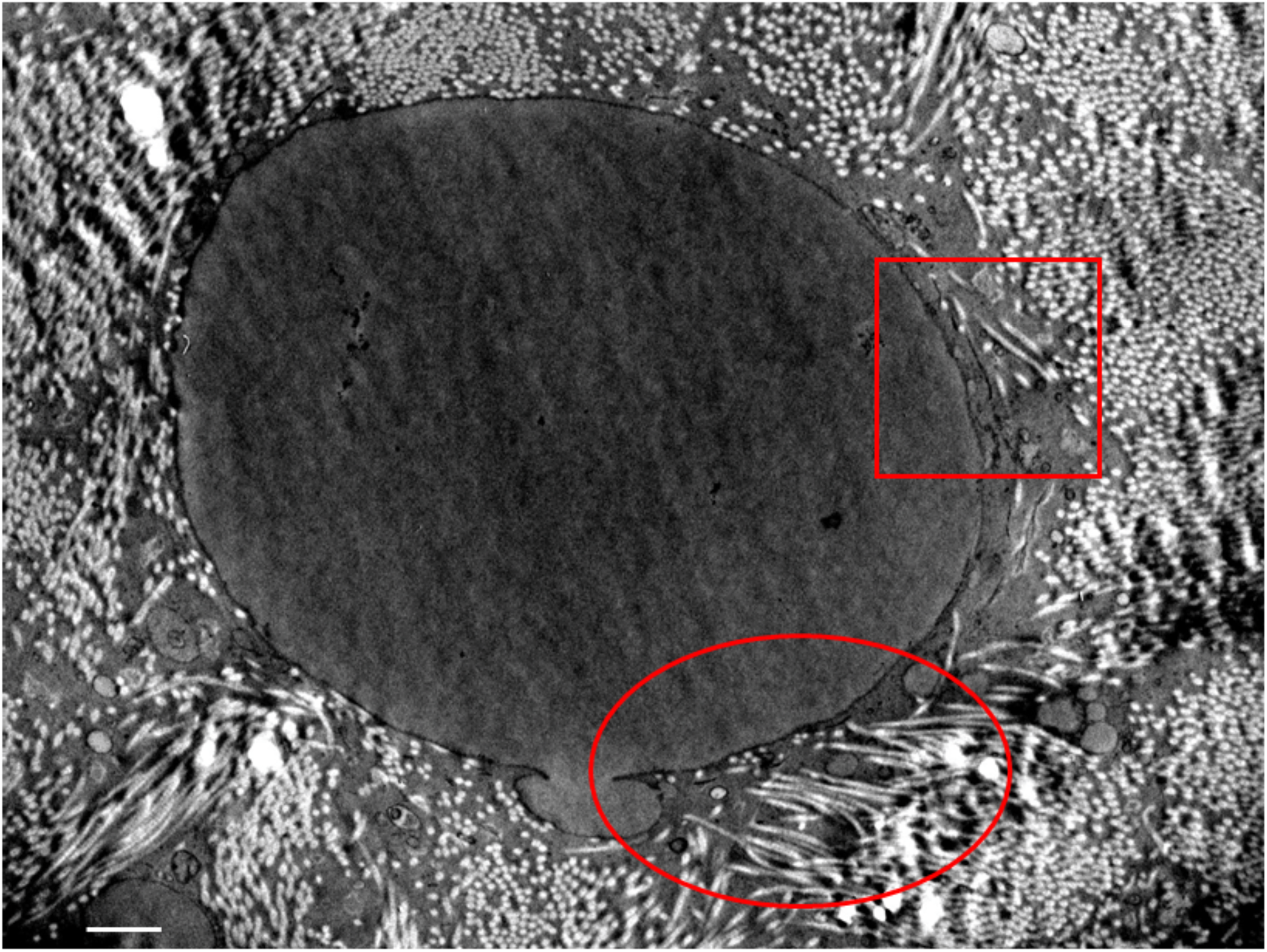
Representative ultrastructure of adipocyte from subcutaneous fat of a patient suffering for Ullrich syndrome (homozygous mutation of COL6A2 gene). Note the abundance of collagen fibrils in thigh connection with the adipocyte surface. Some fibrils are even apparently located into the cytoplasm (squared area, enlarged in Fig 4) and some seems to sprout directly from the cell (circled area). Bar: 2μm

**SFig 7:**
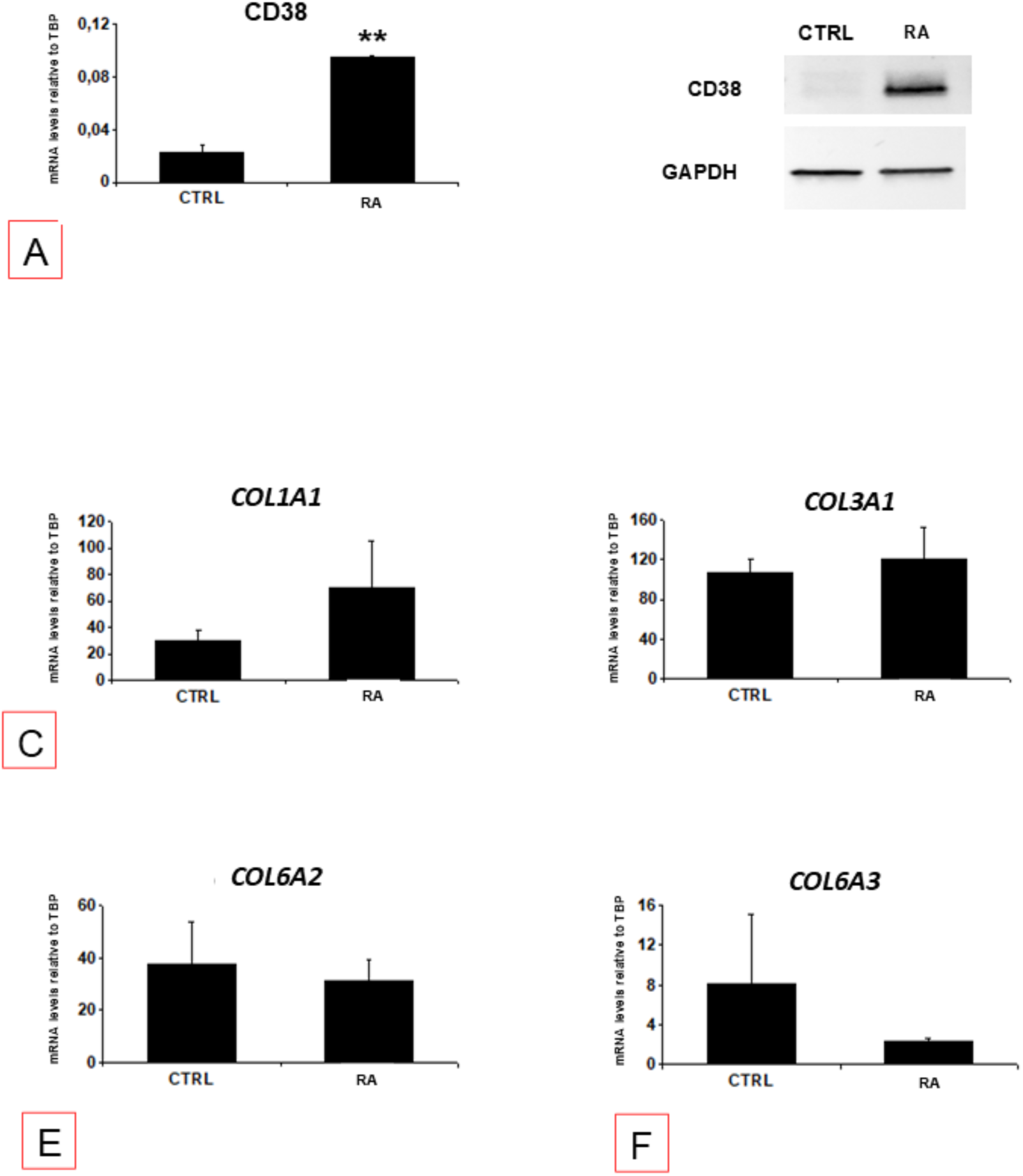
hMADS cells were incubated in the presence or absence of the highly specific inducer of CD38 transcription 1 μM All-trans-Retinoic Acid (RA) for 48 hours. CD38 modulation was analyzed in real time (A) and in western blot (B). Gene expression of COL1A1 (C), COL3A1 (D), COL6A2 and COL6A3 was measured.

**Scheme I:**
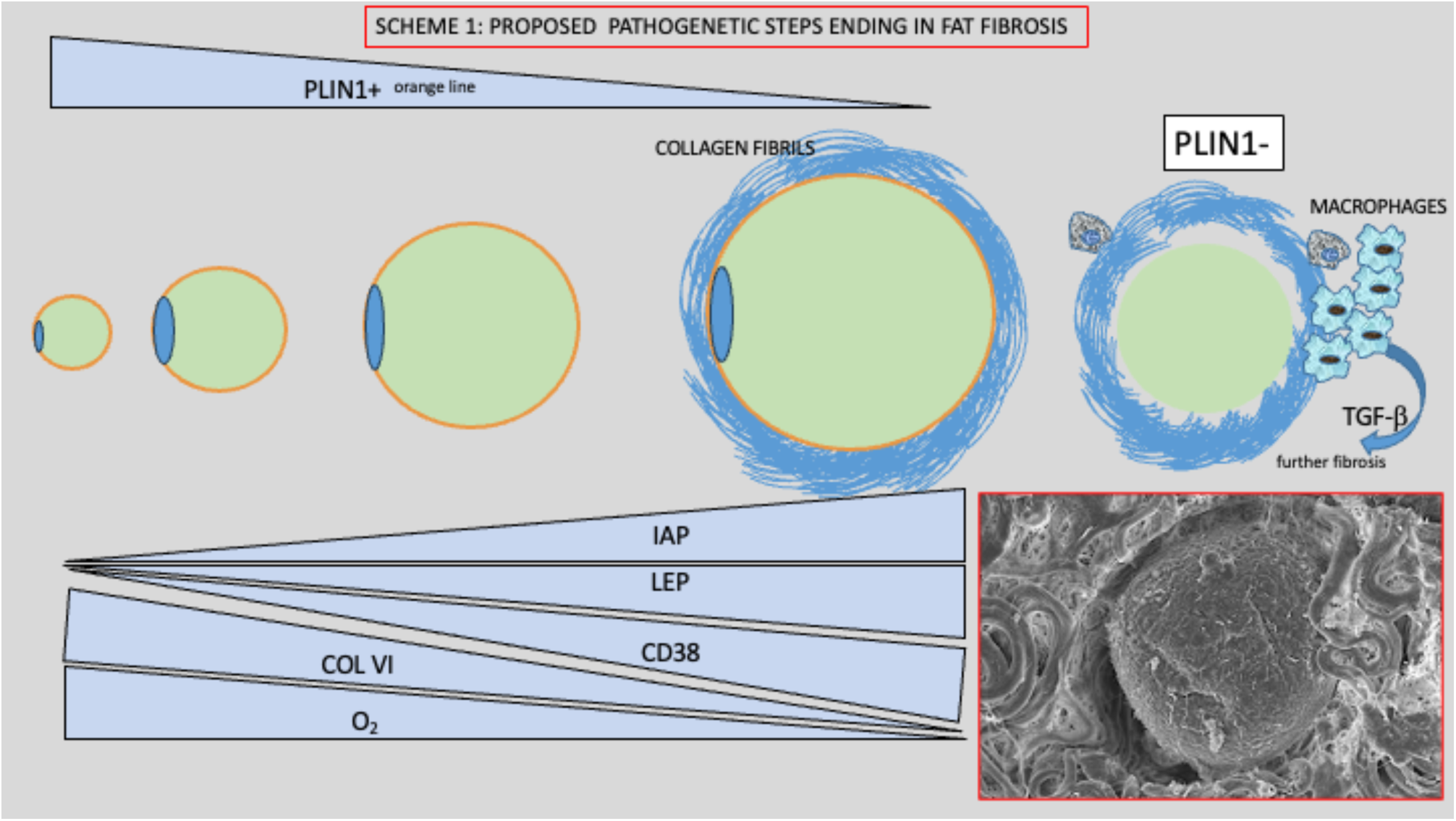
Proposed pathogenetic steps ending in visceral fat fibrosis. With fat accumulation adipocytes undergo a progressive hypertrophy. Hypertrophic adipocytes induce increased intra-abdominal pressure (IAP) with consequent inhibition of COL6 gene expression. COL6 mutations induces high levels of fat fibrosis and could explain the peri cellular fibrosis surrounding hypertrophic adipocytes. Hypoxia due to both hypertrophy and pericellular fibrosis contribute to the stress of hypertrophic adipocytes that lose the perilipin 1 and die. Death of adipocytes induces low grade inflammation mainly due to macrophages. CD38 is highly expressed by hypertrophic adipocytes and by macrophages and stimulates fibrosis trough TGF-β.

